# *Globodera pallida* virulence on major potato resistance has a common genetic basis across Western Europe

**DOI:** 10.64898/2025.12.22.695896

**Authors:** Arno S. Schaveling, Leidy van Rijt, Yoonseon Do, Nike Soffree, Daan Langendoen, Hilde Room, André Machado Bertran, Margien Raven, Sebastiaan P. van Kessel, Evelyn Y.J. van Heese, Stefan J.S. van de Ruitenbeek, Casper C. van Schaik, Sebastian Kiewnick, Geert Smant, Mark G. Sterken

## Abstract

The potato cyst nematode *Globodera pallida* poses a major threat to potato production in Western Europe. Current management strategies largely depend on the use of potato varieties carrying the genetic resistance *GpaV_vrn_*. However, reports from multiple West-European countries indicate a steady rise in virulence against *GpaV_vrn_*-containing potato varieties, raising serious concerns about *G. pallida* control. Although recent studies have resolved the genetic basis of virulence in two Dutch *G. pallida* populations, it remains unclear how conserved this genetic adaptation is in populations from different regions. To investigate this, we first selected eight Dutch *G. pallida* populations on the *GpaV_vrn_*-containing potato variety Seresta and confirmed a previously identified virulence locus. Second, by analysing the allele frequencies of four virulence-associated SNPs in Dutch, British, and French *GpaV_vrn_*-selected *G. pallida* populations, we found that the same allele is consistently selected by *GpaV_vrn_* across Western Europe. Third, we analysed the propagation of eight *G. pallida* populations on 26 *GpaV_vrn_*-containing potato varieties and showed that a population’s allele frequency of a single SNP (T173N) accurately reflects its reproduction on *GpaV_vrn_*. Fourth, we developed an allele-specific quantitative PCR (AS-qPCR) assay to determine a population’s alternative allele frequency (AAF) of T173N and showed that AS-qPCR-based AAFs reliably indicate virulence levels on *GpaV_vrn_* in Dutch and German *G. pallida* populations. Together, these findings suggest that a common allele is consistently selected by *GpaV_vrn_* in populations from different regions across Western Europe. The AS-qPCR assay developed in this study provides a practical tool to estimate *G. pallida* virulence on *GpaV_vrn_* in the field, enabling field-tailored and sustainable resistance management strategies for farmers.

## INTRODUCTION

*Globodera pallida* and *G. rostochiensis* are potato cyst nematodes (PCN) that cause world-wide losses in potato cultivation (Jones et al., 2013). Since the most effective nematicides have been banned (EU, 2009), management strategies primarily rely on crop rotation and the use of PCN-resistant potato varieties. Resistances against PCN in cultivated potato are derived from wild *Solanum* species. *Globodera pallida* resistance in commercial potato varieties is primarily based on *GpaV* locus from *Solanum vernei* (*GpaV_vrn_*; Rouppe van der Voort et al., 2000; Schaveling et al., 2025). However, selection for virulence by *GpaV_vrn_* has led to the rise of resistance-breaking *G. pallida* populations (den Nijs & van Heese, 2019; Mwangi et al., 2019; Niere et al., 2014; Schaveling et al., 2025).

Potatoes and PCN coevolved in the Andes region (Hockland et al., 2012). Here, PCN evolved the ability to hatch upon the perception of potato root exudates. After hatching, juveniles search for and penetrate the potato roots and establish a feeding site (syncytium) that supplies them with nutrients. Meanwhile, potatoes developed the ability to detect and react to invading juveniles by activating defence responses, preventing juveniles from reaching maturity (Goverse & Smant, 2014). Based on its timing, resistances can be classified as male-biased (masculinizing) or blocking. A male-biased resistance, such as *GpaV_vrn_*, acts before the sex determination (Mugniery et al., 2007). It restricts syncytium development, limits a juvenile’s nutrient availability, and thereby promotes male-development. Since male-biased resistances allow avirulent individuals to transmit their alleles to the next generation, this slows down the breakdown of resistance (Schouten, 1994, 1996). In contrast, a blocking resistance, such as *Gpa2*, acts after sex determination, allowing initial syncytium formation before triggering localised necrosis in the cells around the syncytium that eventually starves juveniles (Koropacka, 2010). As female development is highly nutrient-demanding, starvation mainly affects female development (Muller et al., 1981).

PCN are thought to be introduced into Europe through the importation of potatoes in the nineteenth century (Esquibet et al., 2024; Evans et al., 1975; Grenier et al., 2010). Although all West-European *G. pallida* populations originate from the same region in Southern Peru, they are rich in allelic variation (Plantard et al., 2008). Since the introduction of *GpaV_vrn_*-containing potato varieties in the 1990s, these have been extensively used for *G. pallida* control. The deployment of *GpaV_vrn_* exerts positive selection pressure favouring nematodes carrying virulence alleles. Over time, the widespread and repeated use of *GpaV_vrn_* increased the frequency of these alleles leading to breakdown of resistance (Schaveling et al., 2025).

*Globodera pallida* and *G. rostochiensis* are classified as quarantine organisms by the European Commission (EU, 2019), making them subject to strict phytosanitary regulations. Monitoring of PCN involves both voluntary and mandatory sampling of potato fields (EU, 2022; Orlando & Boa, 2023). Upon the extraction and identification of PCN, DNA can be extracted from cysts to determine the *Globodera* species. In cases of suspected virulence, farmers can have the cysts tested on a panel of PCN-resistant potato varieties to determine which variety to grow. However, these variety choice assays are costly and time consuming. Although, long advocated for (Fournet et al., 2018), no rapid, cost-effective and high-throughput test for the detection and quantification of virulence on *GpaV_vrn_* in *G. pallida* has been developed yet.

Recently, a single locus has been associated with virulence on *GpaV_vrn_* in two Dutch *G. pallida* populations and a single gene (*Gp-pat-1*) was found to be directly involved (Schaveling et al., 2025). This gene contained four non-synonymous SNPs that significantly correlated with virulence in two *G. pallida* populations. Building on this work, we confirmed the recently identified virulence locus by bulked segregant analysis on eight *G. pallida* populations that were selected on *GpaV_vrn_*-containing potato variety for four generations. We assessed the allele frequencies of four virulence-associated SNPs in *Gp-pat-1* and found three of these SNPs to be selected across 13 Dutch, French, and British *G. pallida* populations that were all independently selected on potato varieties harbouring *GpaV_vrn_*. For one of these SNPs the frequency of the alternative allele significantly correlated with reproduction on *GpaV_vrn_*. With an allele-specific qPCR (AS-qPCR) we were able to accurately quantify the alternative allele frequency (AAF) of this virulence-associated SNP and identify *GpaV_vrn_*-mediated shifts in the AAF. Finally, AS-qPCR-based AAFs showed to be predictive for virulence on *GpaV_vrn_* in Dutch and German *G. pallida* populations.

## METHODS

### General notes on data analysis

All analyses were conducted in R (version 4.4.1) using Rstudio (version 1.4.1717; R Core Team, 2013; Team, 2015). In R, the *tidyverse* packages, especially *ggplot2* and *dplyr* were used for data processing in general and generation of figures. Other, specific packages used are mentioned at the relevant sections. All R scripts and underlying data are available through git (https://git.wur.nl/published_papers/schaveling_2026_pallida_asqpcr). Sequencing data of the 9 selected and unselected *G. pallida* AMPOP populations was deposited at the European Nucleotide Archive (E-MTAB-16285).

### Selection of *G. pallida* populations on potato variety Seresta and re-sequencing

We generated new sequencing data for nine *G. pallida* populations that were obtained from Dutch potato fields from 2011 until 2015, including the previously described AMPOP10 and eight other populations: AMPOP03, -06, -08, -09, -10, -13, -15, -16, and -19. After initial propagation on Desiree, these populations were selected on the *GpaV_vrn_*-containing potato variety Seresta, as previously described (Schaveling et al., 2025). In short, populations were propagated either for four generations on the resistant potato variety Seresta (S4), or, as a control, for one generation on the susceptible variety Desiree (D; **Supplementary table 1**). Populations resulting from these selection steps were taken as input for DNA isolation and sequencing.

First, DNA extraction and sequencing was performed as previously described (Schaveling et al., 2025). In short, for each S4 and D population DNA of approximately 40 cysts was extracted with phenol-chloroform extraction. DNA was sequenced by BGI Genomics at approximately 200x coverage using DNBSeq with 150bp paired-end reads (Schaveling et al., 2025). For sample AMPOP06D, 27.6 million duplicated reads were removed from FASTQ files with SeqKit (v2.10.0; Shen et al., 2024), after which all data was uploaded to ENA (E-MTAB-16285). This data was supplemented with two pairs of read data for AMPOP02D versus AMPOP02S4 and one pair of AMPOP10D versus AMPOP10S4. These datasets were included in the analysis and previously published by Schaveling et al. (2025). The four independently generated sequencing libraries of AMPOP02 and AMPOP10 were used for method validation.

### Bulk segregant analysis

For analysis of genetic variation the reads were mapped to the Rookmaker genome (PRJEB91928; Schaveling et al., 2025). Variant calling was performed using Bcftools (mpileup & call, v1.14; Danecek et al., 2021) as previously described (Schaveling et al., 2025; https://git.wur.nl/stefan.vanderuitenbeek/dnaseq_variant_calling_snakemake_pipeline). Before bulk segregant analysis, the resulting variant calls were filtered by removing sites with QUAL < 50, variants with a mean read depth outside two standard deviations of the sample average or a minimum read depth of 30, insertions, deletions, and sites with low allelic variation (mean alternative allele frequency ≤ 0.05 or ≥ 0.95). Bulk segregant analysis was performed by conducting a chi-squared test on the read depths for the reference and alternative alleles in the selected and non-selected populations. This is similar to the G-statistic (Magwene et al., 2011) , however as we sequence deep and aim for a high minimum coverage, we calculate the statistic depending on the actual read depths. The resulting p-values were FDR-corrected using the p.adjust function, as we are including variants with linkage disequilibrium.

To identify loci under selection, we counted the number of significant variants in consecutive windows of 10 kb across each scaffold. Identification of bins with a larger number of associated variants was done based on the significance of the numbers based on an exponential distribution. To fit the distribution well, the number of counts per bin was square-root transformed. Bins exceeding the significance threshold (p-value < 0.05) were defined as putative virulence loci.

### Sequencing data of Dutch, British and French populations

Previously, we identified four non-synonymous SNPs inside the coding sequence of a gene involved in virulence on *GpaV_vrn_* (**Supplementary table 2**; Schaveling et al., 2025). To assess how well the AAF of these four SNPs correlate with selection on *GpaV_vrn_*, we used four different DNA sequencing data sets. First, we used the sequencing data of a previous selection experiment with two *G. pallida* populations (AMPOP02 and AMPOP10) selected on the *GpaV_vrn_*-containing Seresta (E-MTAB-15408; Schaveling et al., 2025). Second, we used the sequencing data of the nine *GpaV_vrn_*-selected *G. pallida* populations described above (E-MTAB-16285). Third, we obtained DNA sequencing data of two French *GpaV_vrn_*-selected populations (Saint-Malo and Noirmoutier) from the ENA (PRJEB90550; Lechevalier et al., 2025). Both populations were selected on the *GpaV_vrn_*-containing potato variety Iledher, the *GpaV_spl_*-containing genotype 96D31.51, and the susceptible variety Desiree, *in duplo* for ten consecutive generations. Fourth, we obtained DNA sequencing data of a British *G. pallida* population Newton from the ENA (PRJEB41175; Varypatakis et al., 2020). The Newton population was selected for 12 generations on the *GpaV_vrn_*-containing genotype Sv_11305 (Morag) and the *H3*-containing CPC2802 (Sa_11415) genotype. Mapping and variant calling on the *G. pallida* Rookmaker genome was performed as described above.

### Correlating SNP data with phenotypic data

Two standard PCN resistance tests were conducted in accordance with the EPPO standard protocol (EPPO, 2021). In short, two litre pots with a potato plant were inoculated with approximately 10,000 living larvae. For each population the exact number was determined. At the end of the experiment, cyst numbers were counted per pot. Per pot, the average number of larvae per cyst was determined and multiplied by the total number of cysts in the pot to get the final number of larvae per pot. The reproduction rate was calculated by dividing the final number of larvae (*P_f_*) by the number of inoculated larvae (*P_i_*):

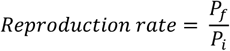

A total of 8 *G. pallida* populations were tested across two tests. Both tests included AMPOP02, -03, -06, - 09, -10 and -16. In addition, test 1 also included AMPOP13 and -15. Test 1 included fourteen potato varieties, that were grouped into clusters based on their resistance level (as determined previously by (Schaveling et al., 2025)). All varieties carry *GpaV_vrn_*. Six varieties were included from the Seresta cluster (Cl_SER_; Axion, Ardeche, Arsenal, VD 07-0289, Innovator, and Seresta) and eight of the Festien cluster (Cl_FES_; Avarna, Avito, Altus, Supporter, Basin Russet, Libero, HZD 06-1249, and Festien). Test 2 included eight varieties from Cl_SER_ (Actaro, Aveka, Novano, Avatar, Simphony, Stratos, Vermont, and Seresta) and six from Cl_FES_ (BMC, Merenco, Saprodi, Sarion, Sereno, and Festien). Each tests contained at least three biological replicates per population (**Supplementary table 3**). The average reproduction on Seresta of both tests was used as an indication of the virulence level on *GpaV_vrn_*.

### An AS-qPCR for the quantification of the alternative allele frequency of T173N

To quantify the AAF of SNP T173N (AAF_T173N_) in *G. pallida* field samples, we decided to develop an allele-specific (AS) qPCR assay. Therefore, we first aimed at specifically amplifying the reference and the alternative alleles. We developed two sets of primers: one standard set and one with a locked nucleic acid (LNA) at the 3’ end. Both allele-specific primers had the SNP at the 3’ end (**Supplementary table 4**). We calculated the qPCR-based AAFs based on the amplification of the alternative allele relative to the total amplification (Germer et al., 2000) according to the formula:

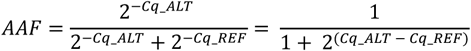

with *Cq_REF* being the Cq-value of the primer pair amplifying the reference allele and *Cq_ALT* being the Cq-value of the primer pair amplifying the alternative allele, reducing the need for housekeeping genes. To obtain DNA for the AS-qPCR, 60-70 cysts were crushed with bead-beating in 150 µL water for 80 seconds at 30 Hz. Lysis was performed with an *in-house* lysis buffer (Holterman et al., 2006). DNA was extracted using the Maxwell_®_ RSC PureFood GMO and Authentication Kit in combination with the Maxwell_®_ RSC instrument (Promega, USA) following the manufacturer’s instructions. This yielded between 4.1 and 11.3 ng/ µL of DNA per sample. AS-qPCRs were performed using iQ Supermix (Bio-Rad, USA; **Supplementary table 5**) on a CFX96 C100 Touch thermal cycler (Bio-Rad, USA; **Supplementary table 6**).

In a first test, these primers were tested on seven *G. pallida* populations, including three *in house* reference populations (D383, Rookmaker and AMPOP10) and four populations that were recently isolated from a field with suspected breakdown of *GpaV_vrn_* resistance. As the standard allele-specific primers gave a better indication of the AAF than the LNA primers, we continued our analysis with the standard primers. In a second test, the standard primers were tested on an additional six recently isolated *G. pallida* field populations.

### Blinded test of the AS-qPCR across four laboratories

To validate the AS-qPCR, we designed a blinded test. This test was conducted across four labs: Dutch General Inspection Service for Agricultural Seeds and Seed Potatoes (NAK), Hilbrands Laboratorium (HLB), Nederlandse Voedsel- en Warenautoriteit (NVWA), Wageningen University and Research (WUR). Each lab tested the same blinded set of 12 *G. pallida* populations. This included 10 *GpaV_vrn_*-unselected and selected populations and two reference populations (D383 and Rookmaker). Each of the labs used its own DNA extraction protocol and qPCR machines. DNA extraction was performed on 20-40 cysts.

The WUR performed the AS-qPCR as described above, but with DNA isolation on 20 cysts, yielding between 101.7 and 281.8 ng/µL of DNA per sample. The NAK crushed the cysts with bead-beating in 100 µL water for 5 minutes at 30 Hz. Lysis was done by adding 300 µL Tissue & Cell Lysis Solution (LGC BioSearch Technologies, USA) and 1 µL proteinase K for 15 minutes at 65 °C shaking at 750 rpm. After 10 minutes on ice, 150 µL pre-cooled MPC Protein Precipitation Solution (LGC BioSearch Technologies, USA) was added and centrifuged at 16,000 rcf for 10 minutes. DNA and RNA was isolated with magnetic beads in a KingFisher Flex system (ThermoFisher Scientific, USA), yielding between 5.7 and 38.7 ng/µL of DNA per sample. AS-qPCRs were performed with PerfeCTa SYBR_®_ Green SuperMix (QuantaBio, USA) on an Applied Biosystems™ 7500 Real-Time PCR system (ThermoFisher Scientific, USA). The HLB crushed the cysts, after which lysis and DNA purification was carried out using the Nexttec Tissue & Cell DNA Isolation Kit (Nexttec, Germany) according to the manufacturer’s instructions. This yielded concentrations ranging from 8.5 to 97.5 ng/µL. AS-qPCRs were performed with SYBR Green PCR Mix (Clear Detections, Netherlands) on a CFX96 thermal cycler (Bio-Rad, USA). The NVWA crushed the cysts with bead-beating in a mix of 50 µL water, 130 µL ATL buffer (Qiagen, Germany) and 20 µL proteinase K for 2 minutes at 30 Hz. Lysis was done at 56 °C for 60 minutes shaking at 800 rpm. DNA extraction was performed with the DNeasy Blood and Tissue Kit (Qiagen, Germany) according to the manufacturers protocol, yielding DNA concentration ranging between 2.4 and 13.8 ng/µL. AS-qPCRs were performed with GoTaq® qPCR Master Mix (Promega, USA) on a CFX Opus Real-Time PCR machine (Bio-Rad, USA).

The methods on the assessment of the reliability of the AS-qPCR assay can be found in the **supplementary text 1**.

### Using Dutch and German *G. pallida* populations to assess the predictive value of the AS-qPCR assay

To assess the predictive power of the AS-qPCR, six Dutch *G. pallida* field populations were tested on five potato varieties, including the susceptible variety Desiree and four *GpaV_vrn_*-containing varieties (Seresta, Festien, Axion, and Supporter) in small container tests as previously described (Schaveling et al., 2025). Cysts were counted at 8-12 weeks after inoculation. Relative susceptibilities were calculated for each resistant variety by dividing the number of cysts on a resistant variety by the number of cysts on a Desiree times 100%.

To assess the value of the AS-qPCR on German *G. pallida* populations, virulence levels of ten German *G. pallida* populations obtained from the Emsland region (Germany) were determined through bioassays routinely used (Price et al., 2024). In short, eye plugs of the susceptible potato variety Desiree or the *GpaV_vrn_*-containing varieties Seresta and Axion were individually planted and inoculated with cysts. White females were counted at six to eight weeks after inoculation. The relative susceptibility (RS) was calculated by dividing the number of white females on resistant varieties by the number of white females on the susceptible control times 100%. Using this type of bioassay allows for fast evaluation of the virulence of a *G. pallida* population (Gartner et al., 2021), however, it can overestimate the virulence compared standardized pot trials as relative susceptibility values can reach up to 100% (Price et al., 2024). All ten German *G. pallida* populations were tested by the AS-qPCR assay. From each population, DNA was extracted from 60 cysts using the MasterPure Complete DNA & RNA Purification Kit (Lucigen, USA) following the manufacturers protocol. The DNA was suspended in 10mM Tris-HCl buffer and DNA concentration was determined by a Qubit_TM_ 4 fluorometer (Life Technologies, Singapore). AS-qPCRs were performed with SSoAdvanced SYBR Green Supermix using a CFX 96 Real-Time PCR machine (Bio-Rad, USA), with 14 to 23 ng template DNA per 20µl reaction in triplicate.

## RESULTS

### Bulk segregant analysis on eight *G. pallida* field populations independently verifies a virulence locus

Previously, we identified a locus that strongly associates with virulence in two *G. pallida* field populations (Schaveling et al., 2025). This locus lies on scaffold 28 of the *G. pallida* Rookmaker genome, with the strongest association between 6,371,140 – 6,682,119 bases. For the identification we conducted whole genome sequencing on each generation of a five-generation selection experiment on two *G. pallida* populations (AMPOP02 and AMPOP10) with one to three replicates per population per generation. The locus was identified based on association of allele-frequency shifts over these five generations. Here we aimed to independently verify that this locus is involved in virulence on *GpaV_vrn_*. To make that feasible, we confirmed in four independent pairs of selected AMPOP02 and AMPOP10 whether bulk-segregant analysis (BSA) on the sequencing pools with only an unselected and a four-generation selected population as input was sufficient to identify the locus in all four tests (**Supplementary figure 1**). From this experiment, we conclude that BSA based on generations of selection is a feasible approach for locus verification.

To independently verify the selected locus, we propagated eight *G. pallida* field populations on the *GpaV_vrn_*-containing variety Seresta for four generations and sequenced the genome pool of starting and final populations (**Figure 1A**). This included *G. pallida* populations AMPOP03, -06, -08, -09, -13, -15, -16, and -19. Next, we performed bulk segregant analyses (BSAs) on each of the eight *G. pallida* populations separately. We identified the previous associated locus in six of the eight populations (**Supplementary figure 2**). To be able to draw an overall conclusion from these separate analyses, we pooled the individual outcomes (number of significantly associated variants per 10kb bin). Over all eight populations we identified a single locus on scaffold 28 from 6.21 – 7.80 Mb (p<0.05; **Figure 1B**). The 95% strongest associated loci (>477 significant SNPs counted over the 8 selected populations per 10kb bin) were located between 6.46 – 6.69 Mb. Since the previously identified virulence locus on scaffold 28 ranged from 6.37-6.68 Mb (Schaveling et al., 2025), we conclude this analysis independently verifies the virulence locus.

**Figure 1.**
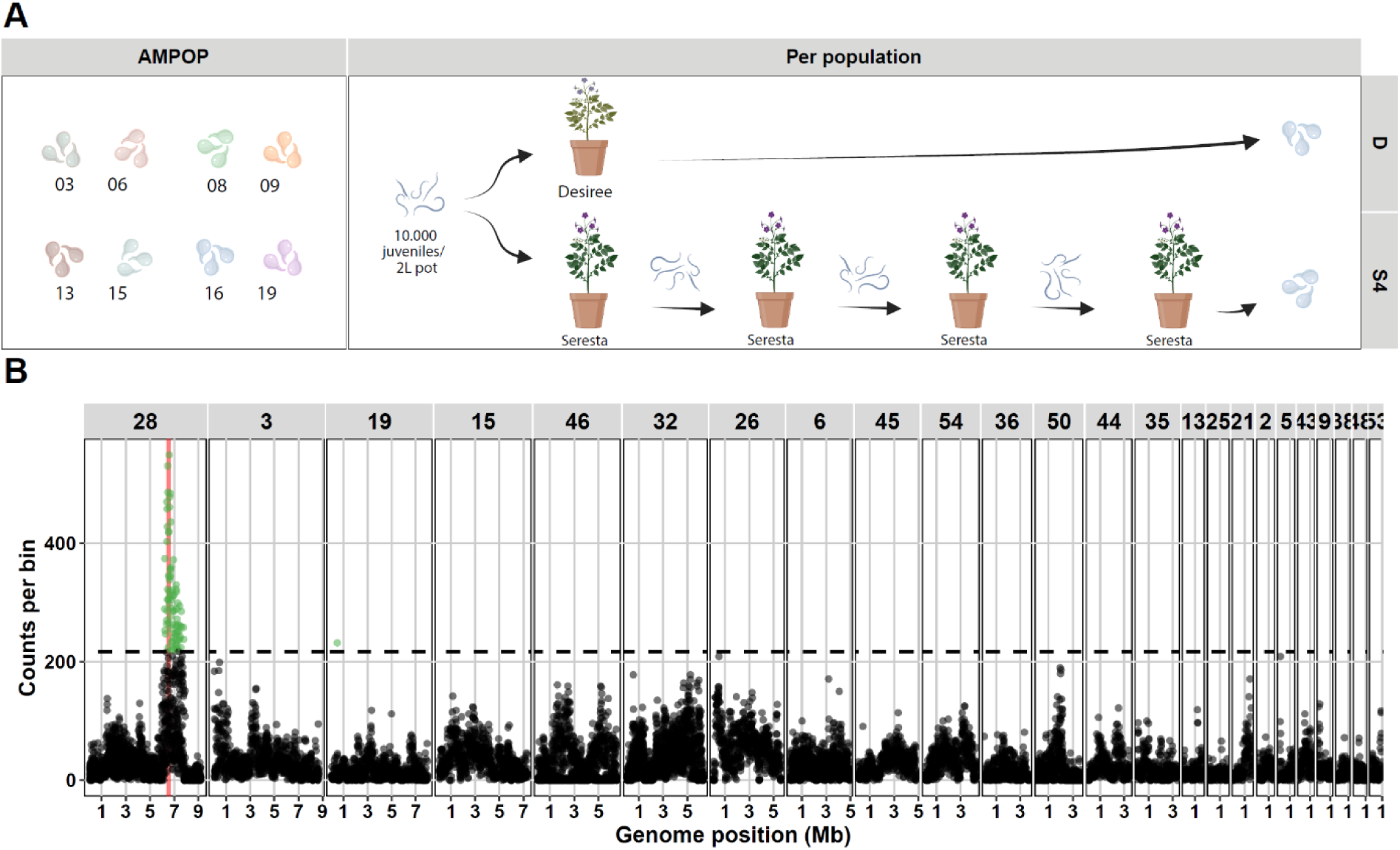
Bulked segregant analysis confirms the previously identified virulence locus. **A** A graphical representation of the eight *G. pallida* AMPOP populations and the selection experiment. Created in BioRender (BioRender.com/v1ycgcu). **B** The number of significant SNPs (FDR<0.05) are plotted per 10kb bin over eight bulk segregant analyses. Bins that are significantly enriched (p<0.05) are coloured in green. The virulence locus is indicated in red.

### Three SNPs are consistently selected by *GpaV_vrn_* across Dutch, British and French *G. pallida* populations

Following verification of the virulence locus, we aimed to investigate whether *G. pallida* virulence on *GpaV_vrn_* has a common genetic basis. Previously, we identified a single gene on the virulence locus, *Gp-pat-1*, as a gene that contributes to the breakdown of *GpaV_vrn_* (Schaveling et al., 2025). Four non-synonymous SNPs (E98V, T173N, L562V, L562S) inside the coding sequence of *Gp-pat-1* correlated with virulence on *GpaV_vrn_* in two *G. pallida* populations (**Supplementary table 2**; Schaveling et al., 2025). To identify allelic variants involved in or tightly linked to virulence on *GpaV_vrn_*, we assessed the alternative allele frequencies (AAFs) of these four SNPs in unselected and *GpaV_vrn_*-selected *G. pallida* populations. First, we assessed the AAFs in two previously selected *G. pallida* populations (AMPOP02 and 10; Schaveling et al., 2025). Over five generations of selection, the four SNPs showed a consistent average increase in AAF of 0.041 per generation (**Figure 2A**). Second, we compared the AAFs of these four SNPs in the nine *G. pallida* populations (AMPOP03, -06, -08, -09, -10, -13, -15, -16, and -19) that we selected on the *GpaV_vrn_*-containing potato variety Seresta for four generations with the unselected populations. For T173N, L562V, and L562S we observed a consistent and significant increase in the AAFs (p-value ≤ 0.00022; **Figure 2B**). This indicates that we have now observed a similar increase in the AAF for three SNPs in a total of 10 genetically unique Dutch *G. pallida* populations.

**Figure 2.**
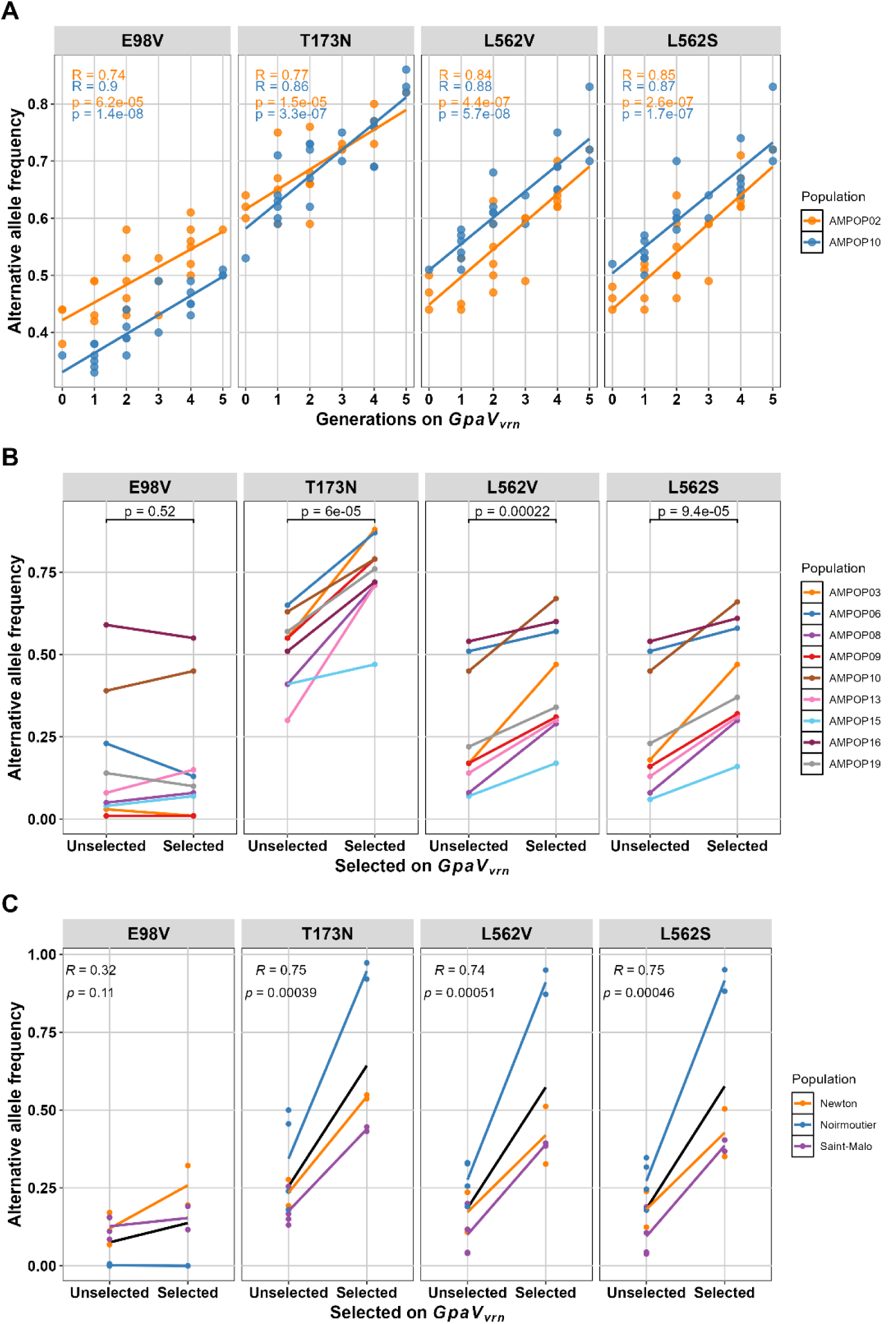
The alternative allele frequencies of four SNPs that are located within the coding sequence of *Gp-pat-1*. **A** The alternative allele frequencies (AAFs) of two *G. pallida* populations selected for five generations on the *GpaV_vrn_*-resistant Seresta potato variety. AAFs show a significant increase over time. Do note that the y-axis is not starting at zero. **B** Nine *G. pallida* field populations were selected for four consecutive generations. SNPs T173N, L562V, and L562S show a significant correlation with selection. Statistics are based on one-sided paired t-tests. **C** Two French *G. pallida* populations (Noirmoutier and Saint-Malo) and an English *G. pallida* population (Newton) show a similar increase in the AAF over time when selected on a *GpaV_vrn_*-resistant potato variety for 10 and 12 generations, respectively. Black lines indicate linear regression models. Statistics in **A** and **C** are based on linear regression models.

Under similar experimental conditions, French and British *G. pallida* populations have shown the potential to break *GpaV_vrn_* resistance as well (Eoche-Bosy et al., 2017; Varypatakis et al., 2019). Two French populations (Saint-Malo and Noirmoutier) were selected on the *GpaV_vrn_*-containing potato variety Iledher for ten generations (Lechevalier et al., 2025) and a British population (Newton) was selected on *GpaV_vrn_*-containing genotype Sv_11305 (Morag) for twelve generations (Varypatakis et al., 2020). To determine the AAFs of the four SNPs in these unselected and *GpaV_vrn_*-selected *G. pallida* populations, we mapped their sequencing data (PRJEB90550 and PRJEB41175) to the *G. pallida* Rookmaker genome. Like the Dutch populations, the French and the British populations showed a significant increase in the AAF for the same three SNPs (**Figure 2C**). In conclusion, three SNPs (T173N, L562V, and L562S) are consistently selected by *GpaV_vrn_* across 13 Dutch, British and French *G. pallida* populations.

### The alternative allele frequency of a single SNP correlates with the reproduction rate on *GpaV_vrn_*

The consistent correlation of three SNPs with selection on *GpaV_vrn_* indicated that these SNPs are either tightly linked to or causal for virulence. To test whether any of our four SNPs gives a robust prediction of virulence, we assessed the reproduction rate (Pf/Pi) of eight *G. pallida* field populations (AMPOP02, -03, - 08, -09, -10, -13, -15, and -16) on 28 potato varieties across two standard PCN resistance tests (EPPO, 2021). These potato varieties were previously grouped into three clusters based on their level of *G. pallida* resistance. These clusters were named after the susceptible potato variety Desiree (Cl_DES_), and the *GpaV_vrn_*-containing potato varieties Seresta (Cl_SER_) and Festien (Cl_FES_; Schaveling et al., 2025). The PCN tests included two Cl_DES_ varieties, thirteen Cl_SER_ varieties and thirteen Cl_FES_ varieties. Since virulence on *GpaV_vrn_* is a recessive trait (Tankam Chedjou et al., 2024), only individuals homozygous for the alternative allele (*aa*) are assumed to be virulent, whereas genotypes *Aa* and *AA* will be avirulent. If we assume that the populations are in Hardy-Weinberg equilibrium, the proportion of *aa* genotypes is expected to equal the square of the AAF. Therefore, to assess a potential quadratic relationship, we plotted the AAF_2_ against the average reproduction rates on Cl_DES_, Cl_SER_, and Cl_FES_. No significant correlations were found between the average reproduction rate on Cl_DES_ varieties and the AAF^2^ of the four SNPs (**Figure 3**). This indicates that the AAFs are not predictive for the reproduction rate on susceptible varieties, which is in line with our expectations. For Cl_SER_, the AAF of T173N significantly correlated with the average reproduction rate on thirteen varieties (p = 0.004). Moreover, when analysed individually, of the six Cl_SER_ potato varieties that were tested by eight *G. pallida* populations, five showed a significant correlation between the AAF of T173N and the reproduction rate (p < 0.05; **Supplementary figure 3A**). For the other three SNPs, no significant correlations with reproduction on Cl_SER_ were found. For Cl_FES_, the average reproduction rate also showed a significant correlation with the AAF of T173N (p = 0.041; **Figure 3**). However, when analysed individually, only in two of the eight Cl_FES_ varieties tested with eight *G. pallida* populations showed a significant correlation between the AAF of T173N and the reproduction rate (p < 0.05; **Supplementary figure 3B**). Together, this data does show that a population’s AAF for T173N gives an accurate prediction of its average virulence level on *GpaV_vrn_*-containing Cl_SER_ and Cl_FES_ varieties, making a promising quantitative genetic marker for virulence.

**Figure 3.**
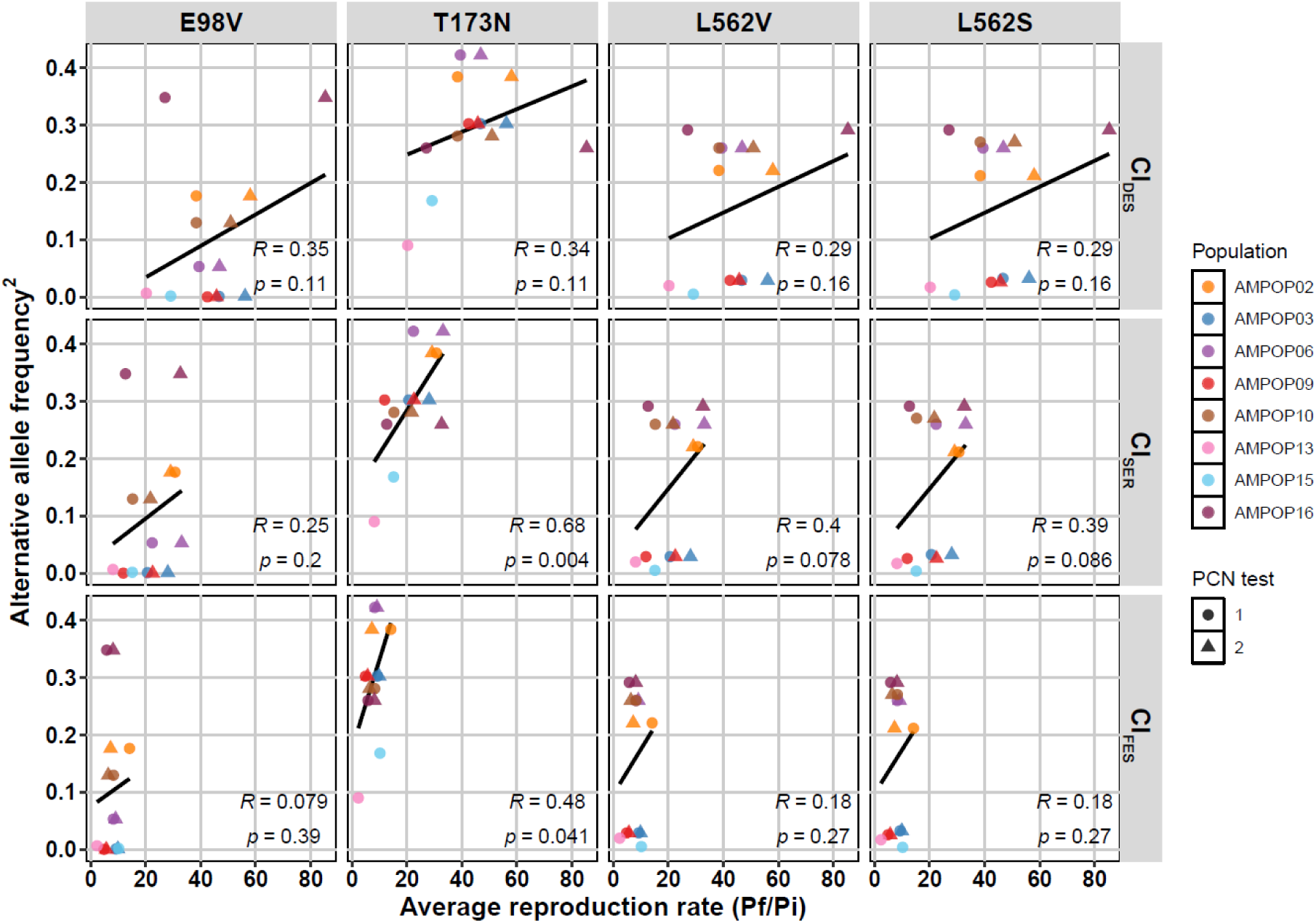
A population’s alternative allele frequency of T173N correlates well with its average reproduction rate on *GpaV_vrn_*. The allele frequencies of eight Dutch *G. pallida* field populations were plotted against their average reproduction rates (Pf/Pi) on two susceptible varieties from the Desiree cluster (Cl_DES_), thirteen *GpaV_vrn_*-containing potato varieties from the Seresta cluster (Cl_SER_) and thirteen *GpaV_vrn_*-containing potato varieties from the Festien cluster (CL_FES_). Significant correlations between the AAF of T173N and the average reproduction on Cl_SER_ and Cl_FES_ indicate that a population’s allele frequency of T173N is a good prediction for its virulence level on *GpaV_vrn_*. Statistics are based on linear regression models that are indicated by black lines.

### Allele-specific qPCR confirms *GpaV_vrn_*-mediated selection of T173N

Now that we have shown that the alternative allele of T173N is consistently selected by *GpaV_vrn_* and the allele frequency in a population correlates with a population’s virulence level on *GpaV_vrn_*, we aimed at developing a PCR-based assay to quantify the AAF of T173N in *G. pallida* field populations. We therefore designed allele-specific primers to specifically amplify the avirulent reference allele and the virulent alternative allele in a qPCR (**Supplementary figure 4a; Supplementary table 4**). First, we tested two variants of the same allele-specific primer sequences: one with a 3’-terminal locked nucleic acid (LNA), and one standard primer. The AAFs as determined by both primer pairs (AAF_qPCR_) significantly correlated with sequencing-based AAF estimates (AAF_seq_; p≤0.0014; **Supplementary figure 4b**). However, the LNA-modified primer overestimated the AAFs, particularly at lower AAFs. In contrast, the standard primers provided an accurate reflection of the sequencing-based AAFs across the tested range. Second, we validated the standard AS-qPCR primers on thirteen *G. pallida* populations, including three previously described populations (D383, Rookmaker and AMPOP10) and ten field populations that we recently isolated from Dutch potato fields. Linear regression models confirmed a strong correlation between the AAF_qPCR_ and the AAF_seq_ (p = 1.5e-10; **Figure 4A**) and the regression coefficients (*β_1_* = 1.00 and *β_0_* = 0.031) indicated that the AS-qPCR assay reliably estimates AAFs across the full AAF range. Third, we assessed the robustness of this test. Across four labs we tested the same set of 22 *G. pallida* populations, including the unselected and Seresta-selected AMPOP02 and AMPOP10 populations (Schaveling et al., 2025), our eight unselected and Seresta-selected AMPOP populations, and reference populations D383 and Rookmaker. The resulting AAF_qPCR_ values of each lab showed highly significant correlations with the sequencing data and with the AAF_qPCR_ values of the other three labs (**Figure 4B**). This indicated that the assay is robust and can accurately quantify the AAF. When we compared the AAF_qPCR_ of the unselected populations with the *GpaV_vrn_*-selected populations, nine out of ten *GpaV_vrn_*-selected populations showed a significant increase in their AAF of T173N as compared to the corresponding unselected population (p < 0.05; **Figure 4C**). Fourth, we tested the AS-qPCR on six *G. pallida* populations previously propagated on potato roots in small container tests. A significant correlation between a population’s average relative susceptibility on *GpaV_vrn_*-containing potato varieties and its AAF_2_ (R=0.83, p=0.021) confirmed that the AS-qPCR is predictive for virulence on *GpaV_vrn_* (**Supplementary figure 5A**). Fifth, we tested the AS-qPCR on ten German *G. pallida* populations. These populations were propagated on the potato varieties Desiree, Seresta, and Axion. A significant correlation between a populations AAF_2_ and the relative susceptibility on Seresta (R=0.61, p=0.031) indicates that the AAF of SNP T173N makes a quantitative genetic marker for virulence on *GpaV_vrn_* in German *G. pallida* populations (**Supplementary figure 5B**). Together, this confirms that *GpaV_vrn_*-mediated selection acts on T173N in each of the tested populations and the AAF of T173N represents a population’s stage of selection.

**Figure 4.**
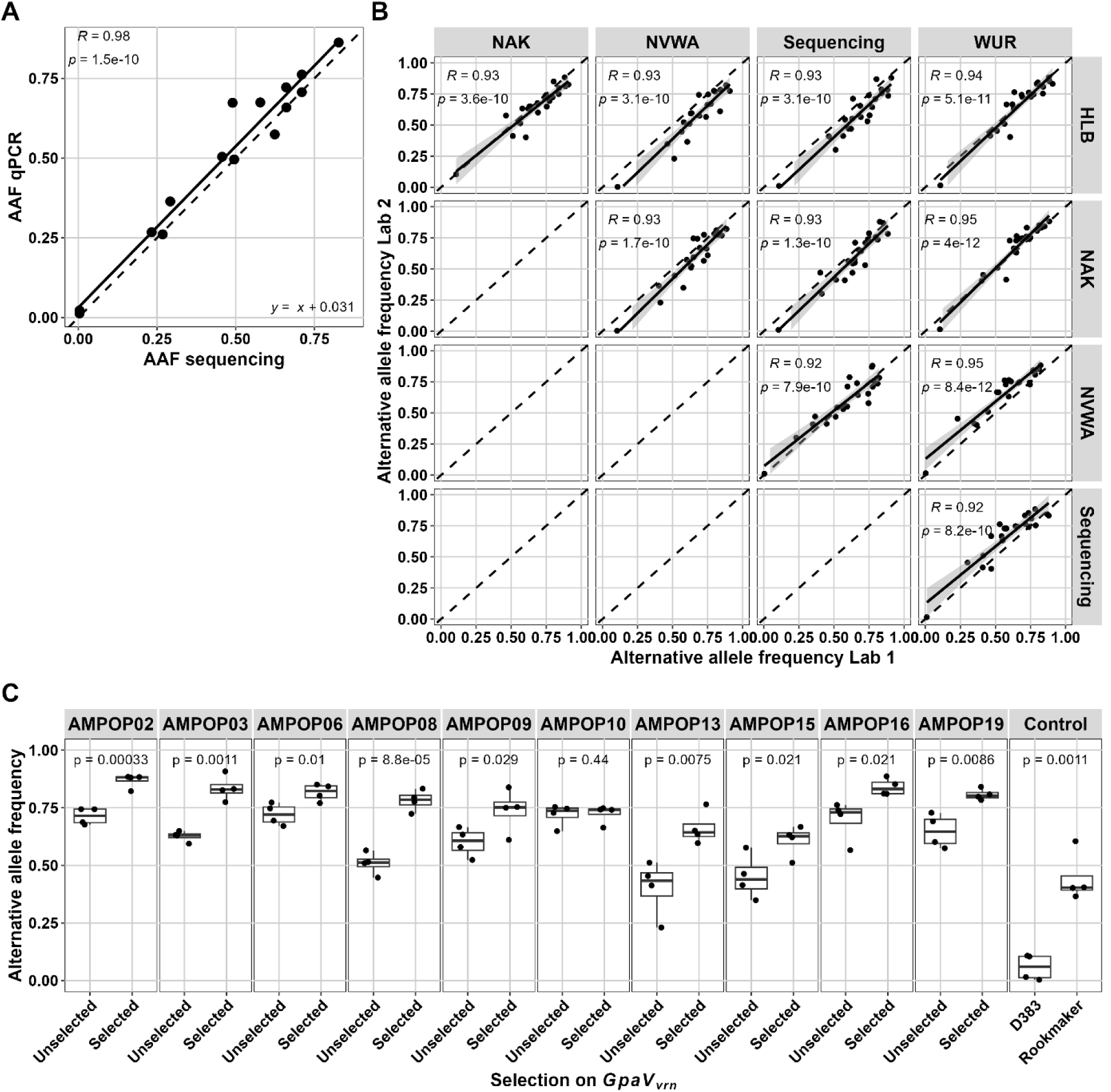
An allele-specific qPCR identifies *GpaV_vrn_*-mediated increases in the alternative allele frequency. **A** The alternative allele frequency (AAF) of SNP T173N in 13 *G. pallida* populations as measured with allele-specific (AS) primers in a qPCR against the AAF according to DNA sequencing. **B** The validation of the AS-qPCR assay at four different labs. AAFs as determined by AS-qPCR by one lab are plotted against the AAFs as determined by qPCR by the other lab or against the AAF as determined by sequencing. Statistics in **A** and **B** are determined by linear regression models. Black dashed lines indicates y=x for visual support. **C** The AAFs of unselected and selected *G. pallida* populations as determined by qPCR in four different labs. Unselected populations were propagated on the susceptible variety Desiree and selected populations were propagated for four rounds on the *GpaV_vrn_*-containing variety Seresta. Statistics are based on one-sided t-tests.

See the **supplementary results** and **Supplementary figure 6** for an assessment on the reliability of this AS-qPCR assay.

## DISCUSSION

Here, we identified a virulence allele in *G. pallida* that is consistently selected across 13 West-European *G. pallida* populations. We showed that its allele frequency (AAF) correlates with a population’s reproduction on *GpaV_vrn_*, and developed an allele-specific qPCR (AS-qPCR) assay predictive for virulence levels of *G. pallida* field populations.

### The allele frequency of a single SNP is a predictor for a population’s level of virulence on *GpaV_vrn_*

Through bulked segregant analyses (BSA) on eight *GpaV_vrn_*-selected and unselected *G. pallida* populations we identified a major virulence locus on scaffold 28 of the Rookmaker genome (**Figure 1B**). This confirms the findings of two previous studies in which *G. pallida* virulence on *GpaV_vrn_* was associated with the same locus on scaffold 28 and the syntenic region on scaffold 2 of the D383 genome (Lechevalier et al., 2025; Schaveling et al., 2025). Collectively, three independent studies, each using a distinct statistical approach (BSA, pairwise *F*_ST_, association analyses) on separate pools of *GpaV_vrn_-*selected *G. pallida* populations, converged on the same virulence locus. In addition to the locus on scaffold 28, we found other loci to be under selection. As these scaffolds are less consistently selected, they may have population-specific contributions to virulence. Given that this is the third independent study to identify the same locus on scaffold 28, strongly supports its central role in virulence on *GpaV_vrn_*.

We then assessed the alternative allele frequencies (AAFs) of four SNPs from *Gp-pat-1*, a gene residing in the associated locus and previously associated with virulence on *GpaV_vrn_* in two *G. pallida* populations (Schaveling et al., 2025). We found that three of the four SNPs were consistently selected by *GpaV_vrn_* across eight other Dutch, one British, and two French *G. pallida* populations (**Figure 2**). This indicates that these SNPs in *Gp-pat-1* are either causal for or tightly linked to virulence on *GpaV_vrn_*. If these SNPs are linked to causal variants, the strength of the correlation between the AAF and virulence will depend on the degree of recombination between the SNP and the causal allelic variant. Across two PCN resistance tests with eight *G. pallida* field populations, we showed that the AAF of SNP T173N significantly correlates with the average reproduction rate on thirteen Seresta-like and thirteen Festien-like potato varieties (**Figure 3**). However, when assessing the correlations for single potato varieties, part of them is insignificant (**Supplementary figure 2**). This likely reflects additional resistances in Cl_FES_ varieties and technical variation inherent to the PCN resistance assay (EPPO, 2021). This is supported by the observation that the only Cl_SER_ variety tested twice (Seresta), showed the strongest correlation (p = 0.00057). Given the technical variation associated with PCN resistance tests, the AAF of T173N may provide a better indication of a population’s average virulence level than individual PCN resistance tests. Therefore, the AAF of T173N represented a reliable indicator of *G. pallida* virulence levels in field populations.

To predict virulence in *G. pallida* field populations, we developed an allele-specific qPCR assay that accurately quantifies the AAF of the T173N SNP and distinguishes populations based on their stage of selection (**Figure 4**). Since the AAF of *G. pallida* field populations was predictive for their propagation on *GpaV_vrn_*-containing potato varieties and the AS-qPCR was able to accurately determine the AAFs, the AS-qPCR assay can serve as a useful tool to predict virulence on *GpaV_vrn_* in field populations.

### Translating allele frequencies to field-tailored advice for potato farmers

With an accurate molecular tool for determining the virulence level of a *G. pallida* population, the major challenge lies in the translation of AS-qPCR data to actionable field predictions. For simplicity, we assumed that our populations are in Hardy-Weinberg equilibrium. We observed significant linear correlations between the AAF^2^ in field populations and their reproduction on *GpaV_vrn_* (**Figure 3; Supplementary figure 6; Supplementary figure 5A**) supporting a quadratic relation between a population’s AAF of T173N and its virulence levels on *GpaV_vrn_*. However, our field populations have been under selection of *GpaV_vrn_*, which has masculinizing activity (Mugniery et al., 2007). When assuming perfect masculinizing resistance, from the second generation onwards, the AAF is expected to show a linear relationship with the number of females and the reproduction rate from the second generation onwards (Tankam Chedjou et al., 2024). However, in the field, perfect masculinizing resistance is not realistic. The ratios in which the allele combinations are present in a population depend on many factors, including the escape rate, the male fraction among virulent individuals and the total fraction of juveniles that is able to reach maturity (Schouten, 1993, 1994; Tankam Chedjou et al., 2024). The relationship between the AAF and the reproduction rate is therefore expected to be in between a linear and a quadratic relationship. Moreover, the effect of the AAF on the reproduction could depend on the infection density and the allele frequency. For example, at low AAFs, having the alternative allele may confer a strong selective advantage, while at higher AAFs intraspecies competition might play a more important role, reducing the benefit of having the virulence allele. Since the overall impact of an infestation depends on both the virulence level of the cysts and their density in the field, translating AS-qPCR-based AAF estimates into actionable field predictions requires integrating AS-qPCR data with cyst densities and phenotypic field data. Future work should therefore establish how infection densities and virulence levels translate into crop loss under field conditions.

Accurate AS-qPCR-based predictions of *G. pallida* virulence levels in the field can be of great help for potato farmers. For example, besides the mandatory PCN sampling (EU, 2022), the majority of Dutch starch potato growers have their fields voluntarily sampled and tested for PCN (Orlando & Boa, 2023). Upon the detection of *G. pallida*, farmers can request variety selection assays to determine which variety to grow based on the *G. pallida* in their fields. However, these assays are costly and time-consuming. Given that soil samples with suspected PCN infestation are already (q)PCR-tested for species and viability determination (Lombard et al., 2024), the infrastructure is there to incorporate a qPCR for virulence determination. In combination with existing qPCR tests, our AS-qPCR test may support service providers in offering farmers informed advice on the sustainable deployment of resistances in the field, tailored to field-specific conditions in a cost-effective manner.

### The same virulence allele is selected by *GpaV_vrn_* across Western Europe

Our analyses showed that the same allele is consistently selected across 13 Dutch, British and French *G. pallida* populations. Moreover, the allele frequency of this same allele correlated with reproduction on *GpaV_vrn_* in Dutch and German *G. pallida* populations. This strongly supports our hypothesis that *G. pallida* virulence on *GpaV_vrn_* has a common genetic basis across Western Europe. The spread of virulence alleles suggests that the emergence of *GpaV_vrn_* resistance-breaking populations can also be expected in West-European countries where no resistance-breaking populations have yet been found. Moreover, since Europe has functioned as a secondary distribution centre for the invasion of PCN into all other continents (Esquibet et al., 2024), the spread of virulence alleles is probably not geographically restricted. This implies that the repeated deployment of *GpaV_vrn_* consistently selects the same virulence locus in *G. pallida*, leading to the rise of similar resistance-breaking *G. pallida* populations irrespective of geography.

## Conclusion

Here, we showed that (1) the same allele is consistently selected by *GpaV_vrn_* in *G. pallida* populations across Western Europe, (2) a population’s AAF of SNP T173N is a good prediction of its reproduction rate on *GpaV_vrn_*, (3) a population’s AAF of T173N can be accurately determined by AS-qPCR, and (4) the AS-qPCR can distinguish different stages of selection. Together, this indicates that virulence on *GpaV_vrn_* has a common genetic basis across Western Europe We conclude that the AS-qPCR-based AAF of SNP T173N accurately reflects a population’s relative virulence level on *GpaV_vrn_*, which makes it the first qPCR-based method for predicting virulence in a parasitic nematode. This assay enables field-specific guidance for sustainable resistance deployment, prolonging the agronomical lifetime of potato varieties, and allows mapping of virulence distribution at different scales. It therefore offers practical value for farmers, service providers, breeding companies, and policymakers.

## Conflict of interest

This research was executed as part of a public/private partnership project funded by the Dutch government including co-financing from several public and private organisations. The authors declare that the research was conducted in the absence of any commercial or financial relationships that could be construed as a potential conflict of interest. After the AS-qPCR results were unblinded, one of the private partners involved in the validation, HLB, began considering offering the assay as a service to growers.

## Author contributions

ASS, GS, and MGS designed the experiments. DL and HR conducted the selection experiments. CCvS and NS assisted in DNA isolation. SJSR assisted in processing and archiving the sequencing data. ASS and MGS conducted the BSA. The qPCR assay was developed by ASS and YD. Validations on Dutch populations were conducted by ASS, AMB, MR, SPvK, and EYJvH. Choice variety assays were performed by AMB and MR. LvR and SK validated the qPCR on German populations. ASS, GS, and MGS wrote the paper with input from all other co-authors.

## Funding

This research was conducted in the framework of the PPS projects PALLIFIT and PALLIGEN funded by PPS subsidies from the Dutch Ministry of Agriculture, Nature and Food Quality and Topsector T&U (KV1604-022/TU-16004 and LWV22225/TU202202, respectively). MGS was supported by NWO domain Applied and Engineering Sciences VENI grant (17282) and NWO domain Applied and Engineering Sciences VIDI grant (21240). LvR received funding from the Agency for renewable resources, project “ASPARA” (FKZ 2222NR093B).

## Acknowledgements

The authors want to thank the private partners in the PALLIFIT and PALLIGEN project for their support and their constructive attitude towards this project. We want to thank Paul Heeres for his role in selecting the AMPOPs on Seresta. We want to thank Kostas Gaitanis for the first attempt to design allele specific qPCR primers. We would like to thank Hemanth Konigopal for technical support in generating high quality template DNA from German *G. pallida* populations. We want to thank three growers for making their fields available to us to collect cysts to use in the validation. We want to thank the Nem-Emerge consortium for discussions regarding the paper.

## Data availability

All scripts and underlying datasets are available through gitlab (https://git.wur.nl/published_papers/schaveling_2026_pallida_asqpcr). The data of presented experiments has been included in supplementary files. The DNA sequencing data of the selection experiments is deposited at BioStudies (E-MTAB-16285).

## SUPPLEMENTARIES

### Supplementary text 1 METHODS

#### Statistical analysis on the reliability of the AS-qPCR assay

To assess how well the AS-qPCR-based alternative allele frequency (AAF) matched the sequencing-based AAF, we used linear regression models that followed the formula:

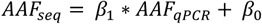

where *AAF_seq_* is the AAF determined by sequencing, *AAF_qPCR_* is the AAF determined by qPCR, *β_1_* is the slope of the model and *β_0_* is the intercept.

To assess the reliability of the qPCR assay, we assumed that the population is in Hardy-Weinberg equilibrium, so we can calculate the standard error (SE) with the formula:

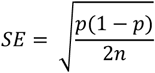

and the effective population size with the formula:

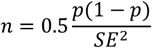

in which *p* is the allele frequency and *n* being the number of individuals (Nei, 1978). The nonlinear least squares model followed the formula:

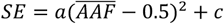

in which *AAF* is the mean *AAF* of a sample as determined by AS-qPCR. The significance of the curvature parameter (*a*) was tested by calculating a one-sided p-value based on the t-statistic (estimate / standard error), using the residual degrees of freedom from the model fit.

### AS-qPCR on mixed populations

To assess the *G. pallida* specificity of the AS-qPCR assay, we performed the assay on *G. rostochiensis* DNA. To assess potential overestimation of the AAF in a mixed population of *G. pallida* and *G. rostochiensis*, we used the formula:

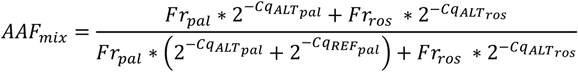

in which *AAF_mix_* is the AAF in a mixed population, *Fr_pal_* is the fraction of *G. pallida* in a population and *Fr_ros_* the fraction of *G. rostochiensis*.

## RESULTS

### AS-qPCR reliability depends on the community composition, AAF and cyst number

Our AS-qPCR assay showed both accuracy and robustness. Next, we aimed at determining its reliability. Our previous AS-qPCR tests used 20 cysts per sample. However, in field samples, cyst numbers are often limited. Since no sample perfectly represents the population in the field, the reliability of the test depends on the number of cysts that can be obtained from a soil sample. To assess the reliability of the assay on low cyst numbers, we calculated the theoretical standard error (SE) based on the AAF and the number of individuals tested (*n*). Although each egg within a cyst is genetically unique, they are not genetically independent, due to shared parentage. Therefore, we considered each cyst to be a single individual. Under a worst-case scenario, where the AAF is 0.5, the theoretical SE is 0.079 [sqrt(0.5_2_/40)]. In practice, however, the SE observed in our AS-qPCR data was lower, ranging from 0.012 to 0.061 (mean = 0.030; **Supplementary figure 6**). This was expected, as not all tests represent worst-case scenarios and because cyst nematode females are polyandrous and can mate with multiple males (Green et al., 1970; Triantaphyllou & Esbenshade, 1990). Therefore, each cyst contains more genetic diversity than the theoretical model assumes, resulting in a larger effective population size in practice. For each sample, we calculated the effective population size based on the observed mean AAF and the SE. This yields an average effective population size of 187 individuals, which is over nine times higher than the assumed population size of 20. To estimate the SE based on the number of cysts, one could use nine times the number of cysts as the number of individuals used (*n*). When the theoretical SE is too high, reducing the SE by 50% requires a fourfold (2_2_) increase in cyst numbers. Conversely, halving the number of cysts increases the SE by a factor 1.41 (√2).

Based on the formula used for calculating the SE, we hypothesised that the SE would be AAF dependent. Fitting a quadratic model with a fixed vertex at AAF = 0.5 confirmed this hypothesis (p = 0.016; **Supplementary figure 6**), indicating that AAF_qPCR_ estimates are least precise for AAFs around 0.5 and more reliable at high and low AAFs. Ultimately, the minimum cyst number required for a reliable AS-qPCR assay depends on what someone considers an acceptable level of precision. Notably, the standard error increases as fewer cysts are used, reducing the reliability of the allele frequency estimate.

### Running the AS-qPCR on mixed populations

To assess the specificity of the assay, we included *G. rostochiensis* samples as a negative control. The primers targeting the reference allele did not show any amplification on *G. rostochiensis* (Cq > 35.0), which is in line with expectations since the reference allele is not present in *G. rostochiensis*. However, *s*ince the primer targeting the alternative allele has just one mismatch on *Gr-pat-1*, the primers targeting the alternative allele did show minor amplification (Cq ∼ 31.3). This may cause a slight overestimation of the AAF in mixed populations. Based on our data, this may lead up to an overestimation by a fraction 0.0024 in case of a populations consisting for 50% of *G. pallida* and 50% *G. rostochiensis.* This overestimation increases when *G. rostochiensis* makes up a larger fraction of the population. However, as the maximum potential overestimation is considerably smaller than the observed standard errors of the AS-qPCR, the potential overestimation of the AAF by *G. rostochiensis* can be ignored.

### Supplementary tables

**Supplementary table 1**

An overview of the DNA sequenced samples from the selection experiments used for bulk segregant analysis. The associated sequence files are listed, the population the sequencing belongs to (Population), the variety the population was propagated on, the total number of reads generated (in millions), the total number of reads that mapped (in millions), the median coverage on the Rookmaker genome, and comments.

**Supplementary table 2**

An overview of the four SNPs located inside the coding sequence of *Gp-pat-1* that were previously associated with virulence on *GpaV_vrn_*.

**Supplementary table 3**

Data of the two standardized PCN resistance tests. A total of eight *G. pallida* populations were tested on 28 potato varieties.

**Supplementary table 4**

A list of the oligonucleotides used in the AS-qPCR.

**Supplementary table 5**

The reaction setup of the AS-qPCR.

**Supplementary table 6**

The details of the AS-qPCR program.

---

**Supplementary figure 1.**
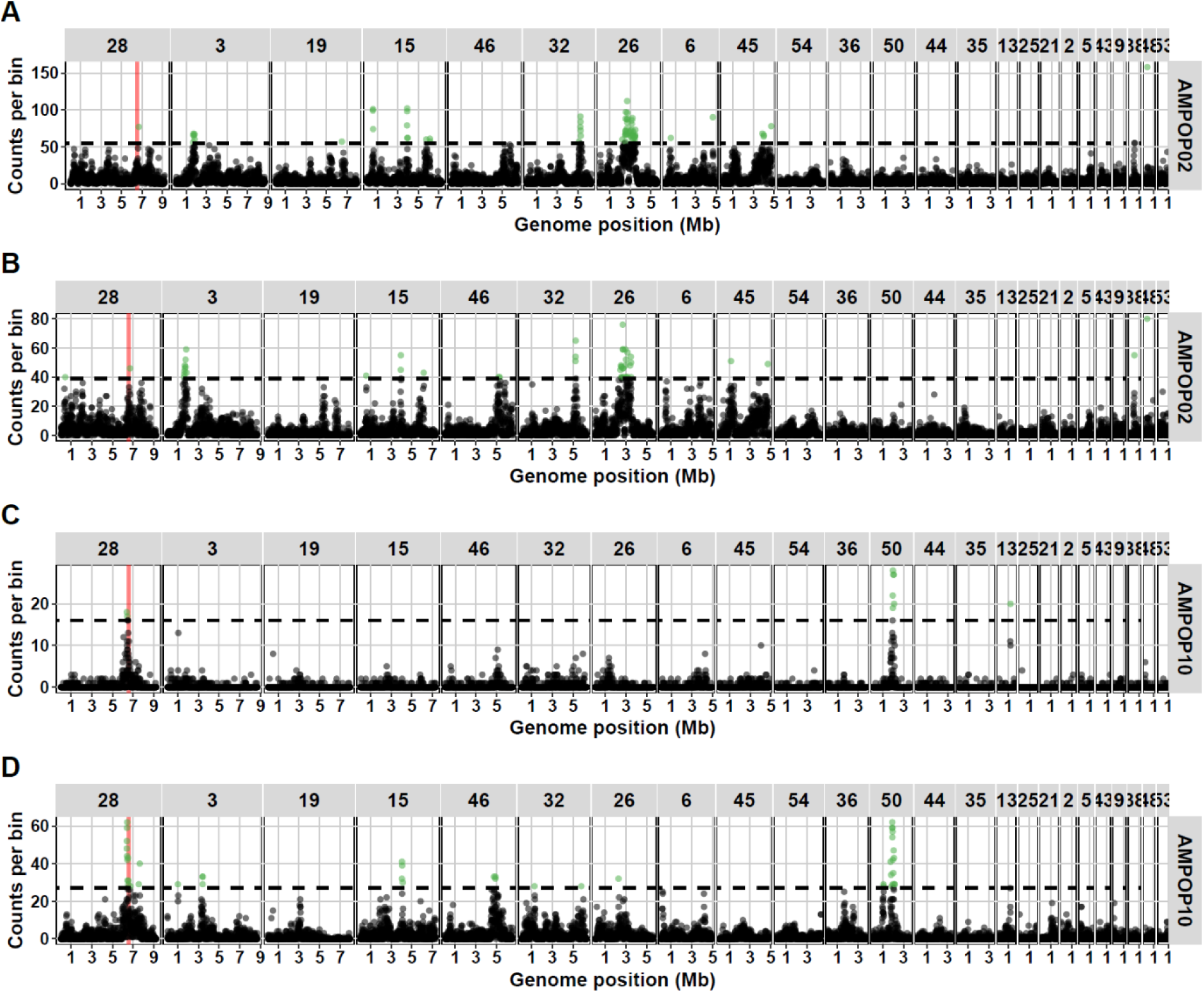
Pairwise bulk segregant analyses between two unselected and *GpaV_vrn_*-selected populations. The number of significant SNPs (FDR<0.05) are plotted per 10kb bin over eight bulk segregant analyses. Bins that are significantly enriched (p<0.05) are coloured in green. The virulence locus is indicated in red.

**Supplementary figure 2.**
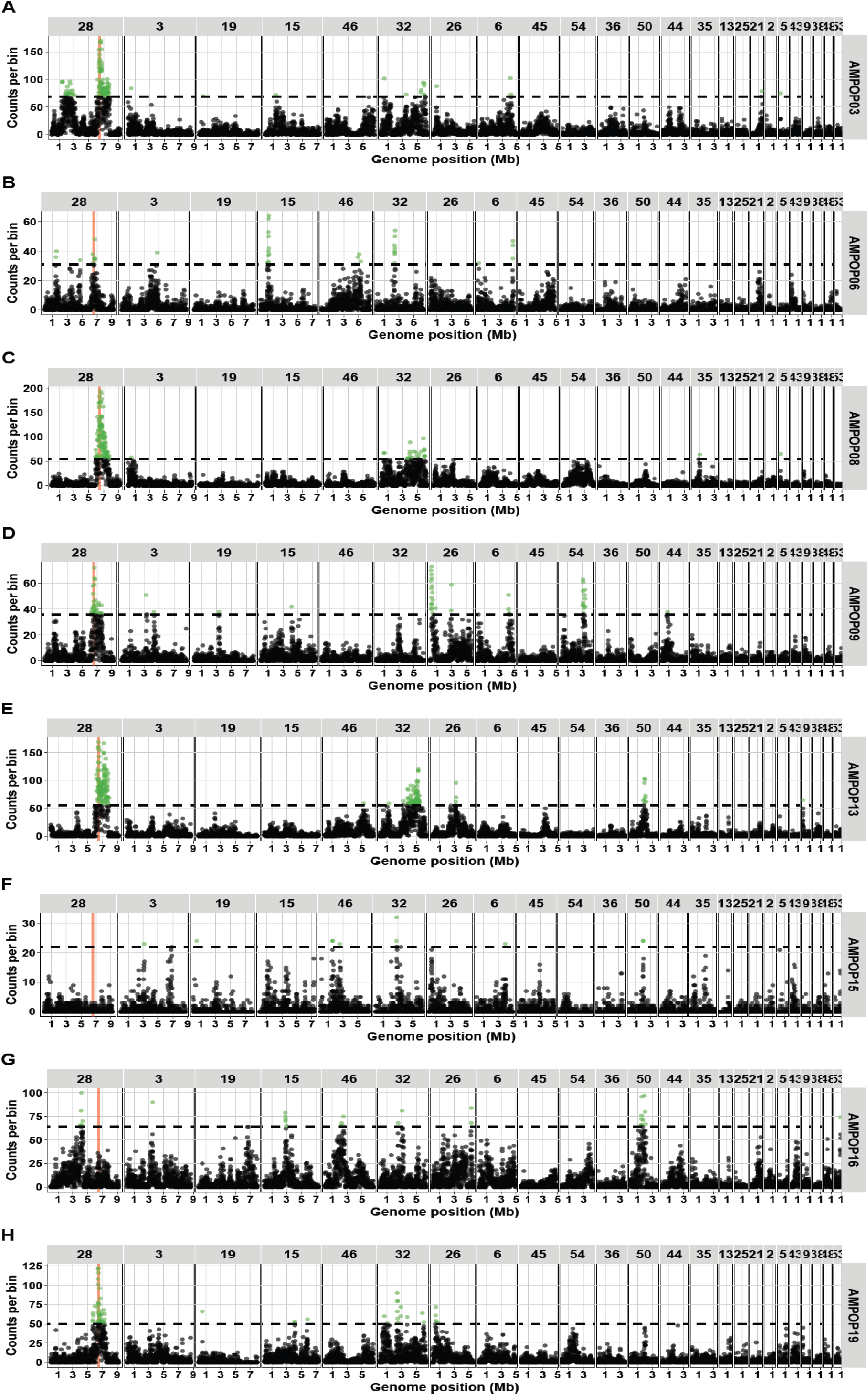
Pairwise bulk segregant analyses between eight unselected and *GpaV_vrn_*-selected populations. The number of significant SNPs (FDR<0.05) are plotted per 10kb bin over eight bulk segregant analyses. Bins that are significantly enriched (p<0.05) are coloured in green. The virulence locus is indicated in red.

**Supplementary figure 3.**
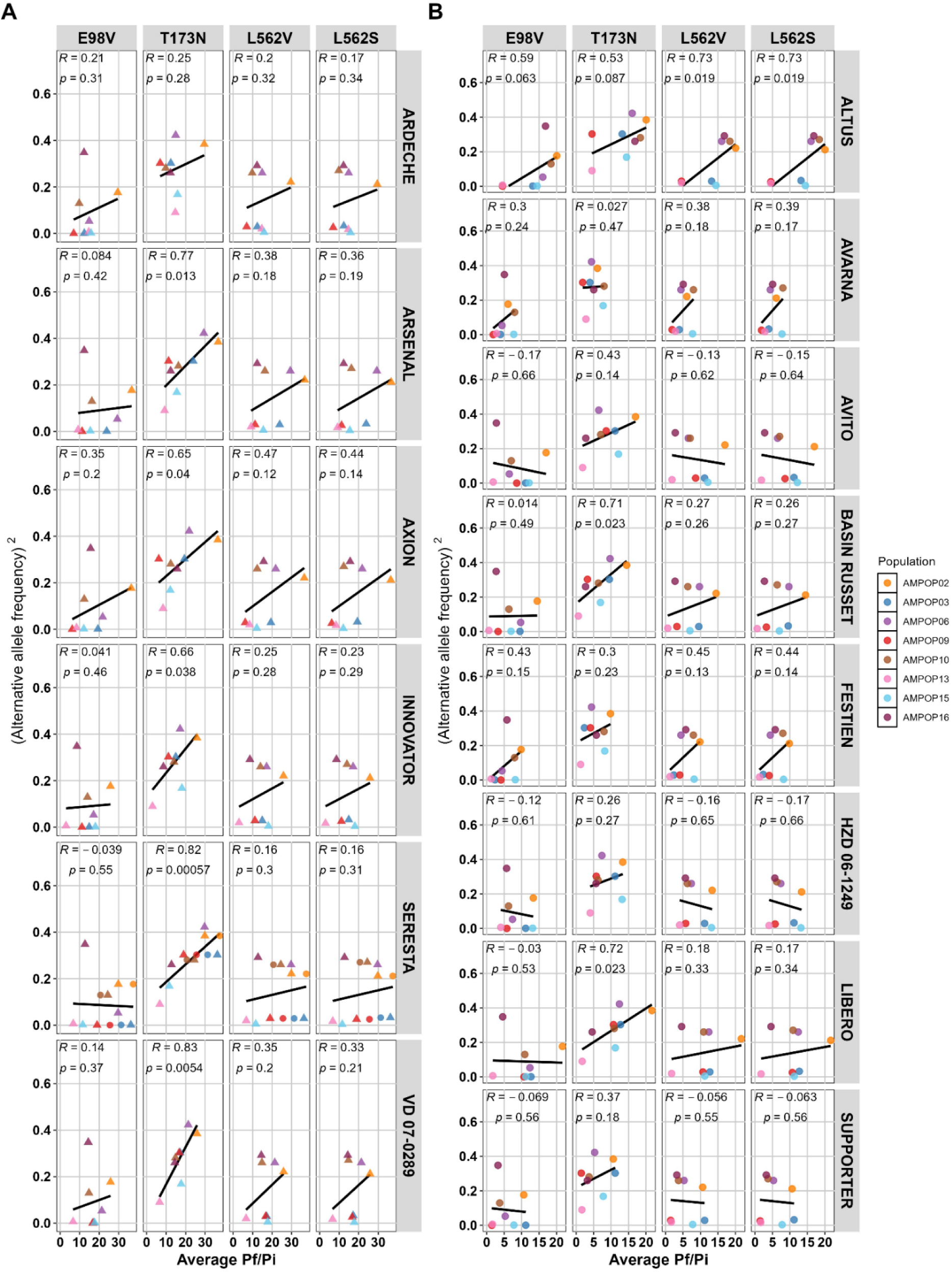
The alternative allele frequencies (AAF) of eight Dutch *G. pallida* field populations were plotted against their corresponding reproduction rates (Pf/Pi) on **A** six different *GpaV_vrn_*-containing potato varieties from the Seresta cluster and **B** eight *GpaV_vrn_*-containing varieties from the Festien cluster. The AAF of T173N significantly correlates with virulence on four of the six Cl_SER_ varieties and 2 of the 8 Cl_FES_ varieties, indicating that the allele frequency of T173N is a good indication for virulence on *GpaV_vrn_*. Statistics are based on linear regression models that are indicated by black lines.

**Supplementary figure 4.**
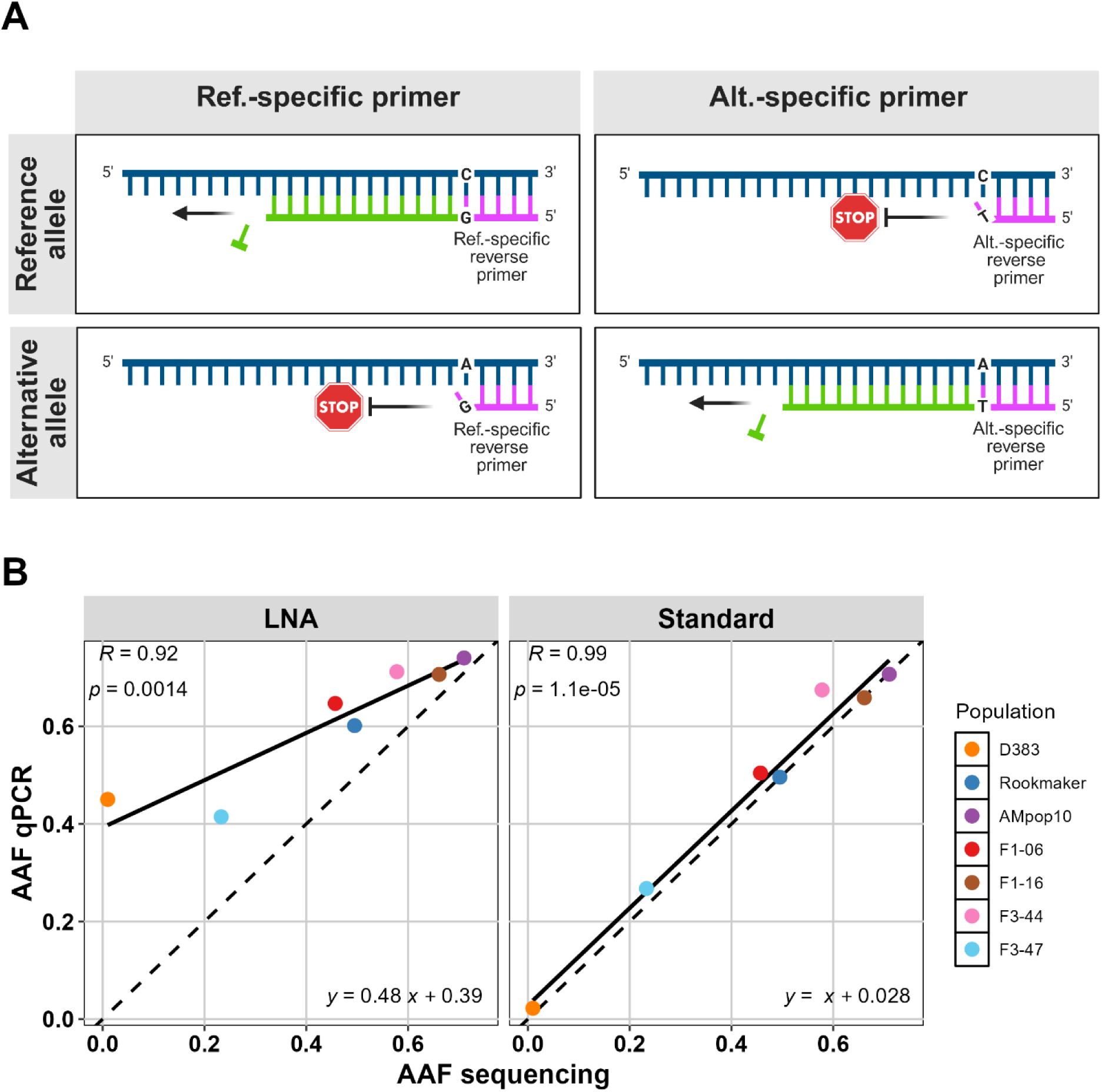
A. An overview of the allele-specific (AS) qPCR design. Template DNA is depicted in blue, primers are depicted in magenta, and DNA amplification in green. Created in BioRender (https://BioRender.com/09y6291). **B** Comparison of two sets of allele-specific qPCR primers: one using LNA-modified primers, with a 3’-terminal locked nucleotide, and one using standard primers. While alternative allele frequency (AAF) estimates from LNA primers significantly correlate with sequencing-based AAFs, they tend to overestimate the AAF in a population, particularly at lower AAFs. In contrast, standard qPCR primers provide a more accurate reflection of the sequencing-based AAF across the tested range. Dashed lines indicate y=x and are added for visual purposes. Statistics are based on linear regression models.

**Supplementary figure 5.**
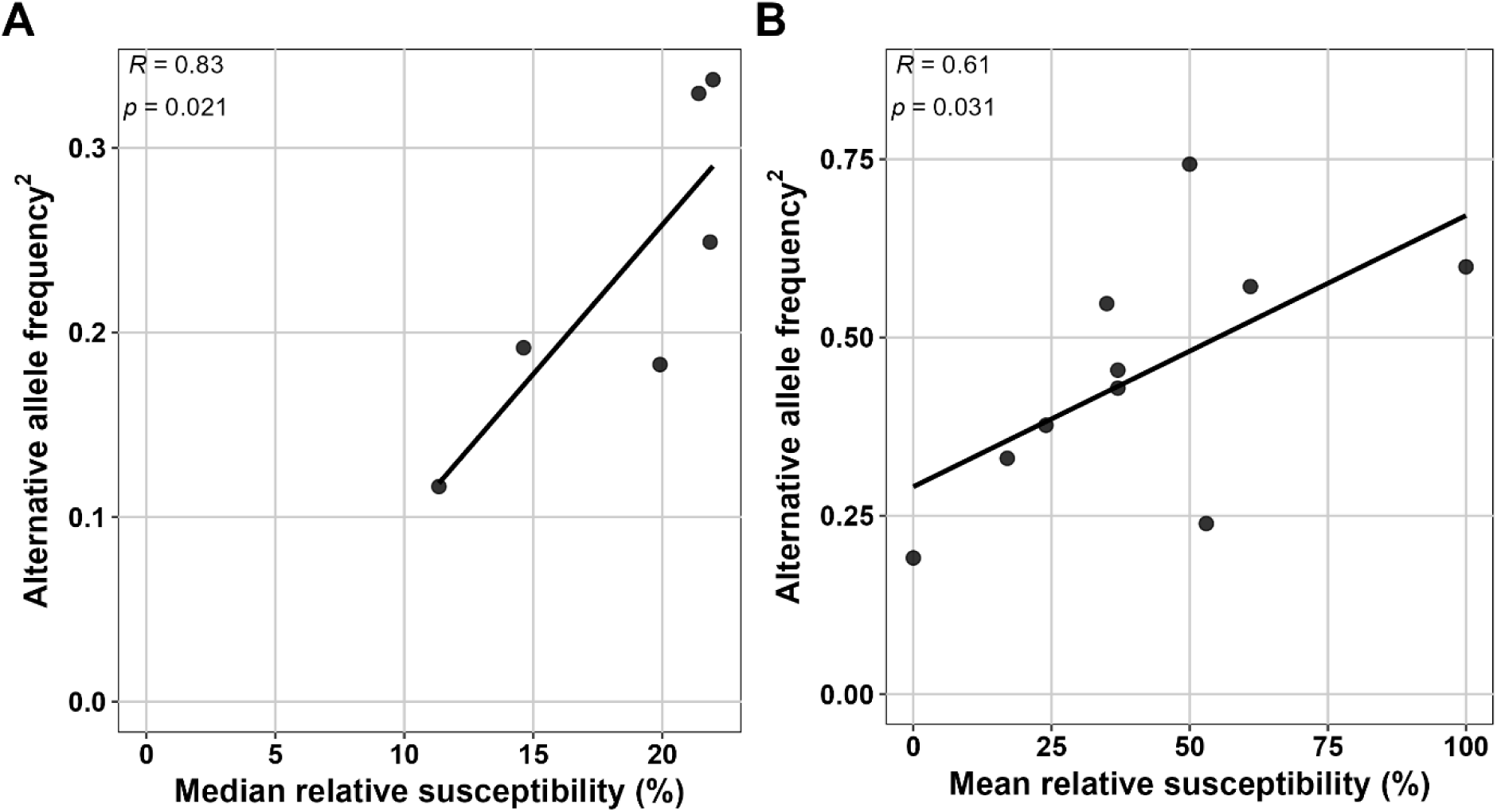
(**A**) The allele frequencies of six Dutch *Globodera pallida* field populations were plotted against their median relative susceptibility (%) on four *GpaV_vrn_*-containing potato varieties. Virulence levels were tested in a small container test and taken relative to the susceptible potato variety Desiree. (**B**) The allele frequencies of ten German *G. pallida* field populations were plotted against their mean relative susceptibility (%) on the *GpaV_vrn_*-containing potato varieties Seresta and Axion. Virulence levels are based on counts of white females in small container tests and taken relative to the susceptible potato variety Desiree. The significant correlations between the AAFs of SNP T173N and the average relative susceptibilities indicates that a population’s allele frequency of T173N is a good prediction for its virulence level on *GpaV_vrn_*. Statistics are based on a linear regression model that is indicated by the black line.

**Supplementary figure 6.**
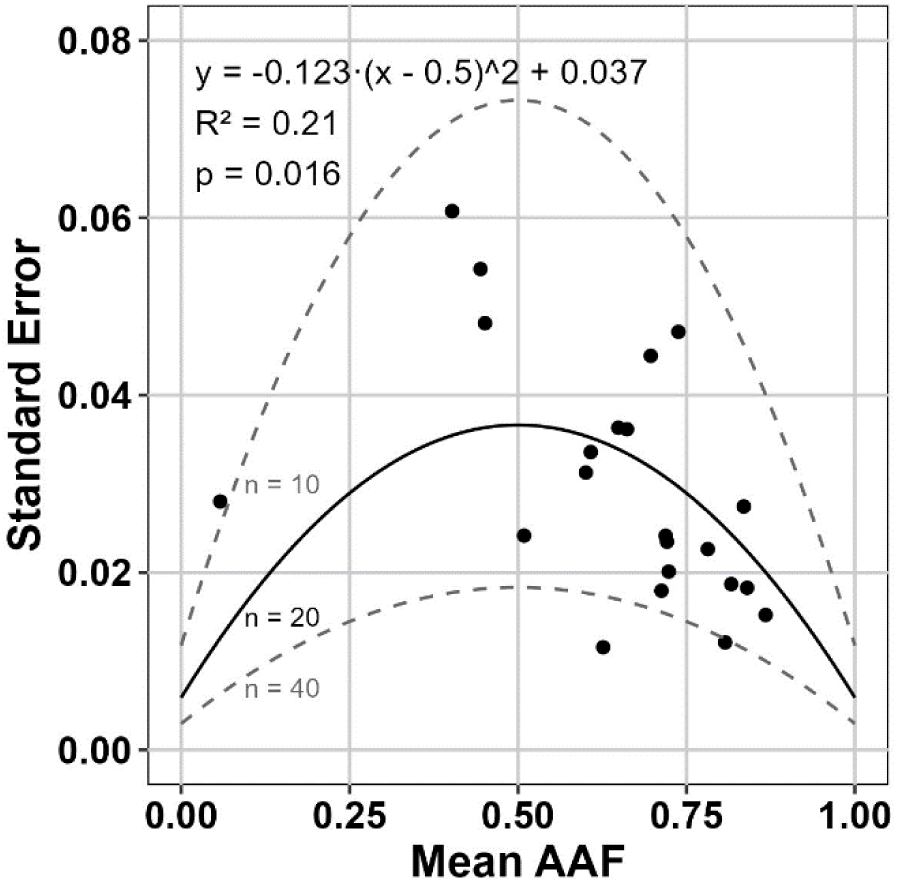
The observed quadratic relation between the standard error and the mean alternative allele frequency (AAF) indicates that the reliability of the assay depends on the AAF of the population (Solid black line). The dashed grey lines indicate the expected standard errors when using halve (n=10) or double (n=40) as many cysts in the AS-qPCR.

## Notes

https://git.wur.nl/published_papers/schaveling_2026_pallida_asqpcr

https://www.ebi.ac.uk/biostudies/ArrayExpress/studies/E-MTAB-16285

