## Supplementary figures and images for "*Globodera pallida* virulence on major potato resistance has a common genetic basis across Western Europe"

### Supplementary figure 1

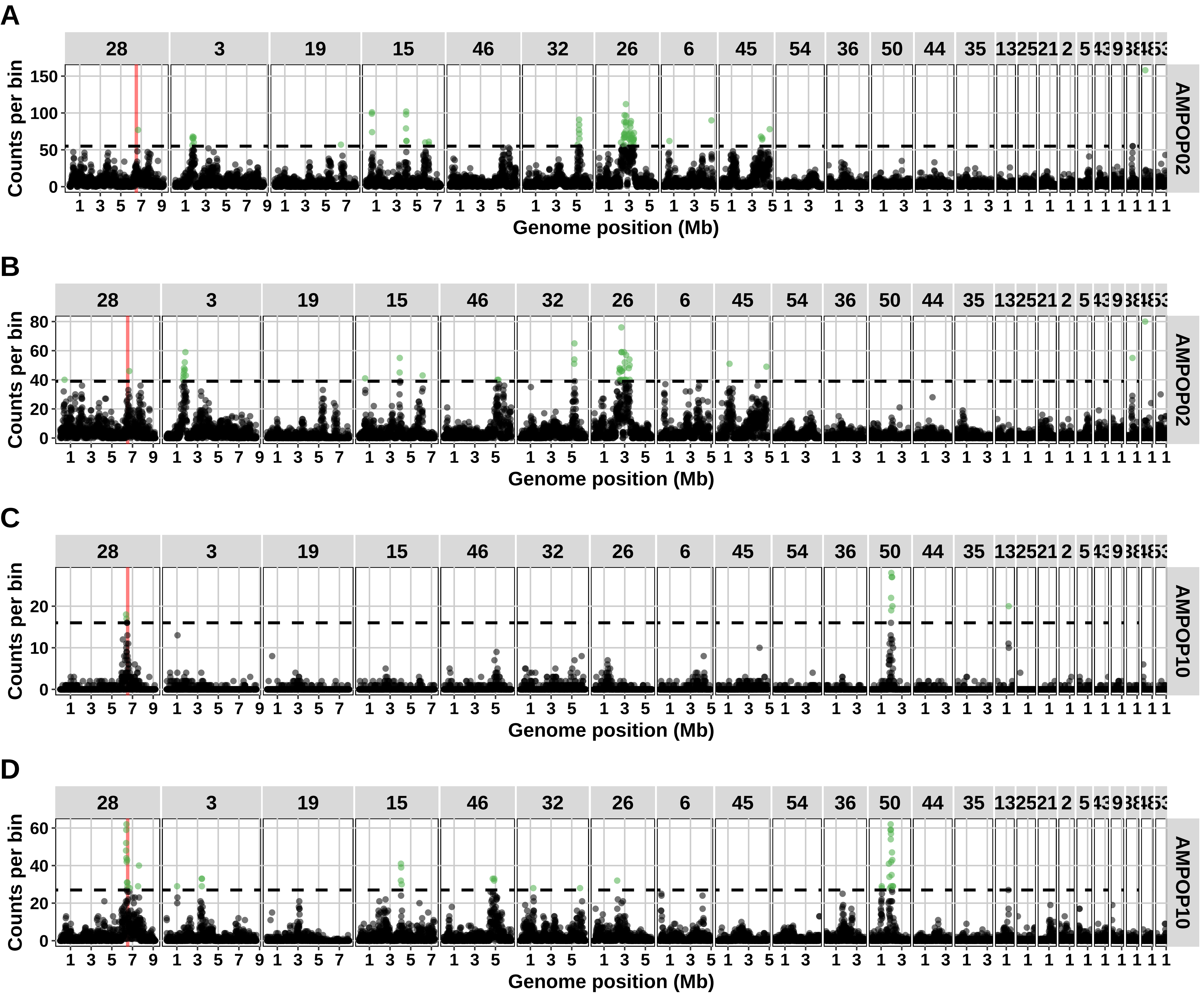

### Supplementary figure 2

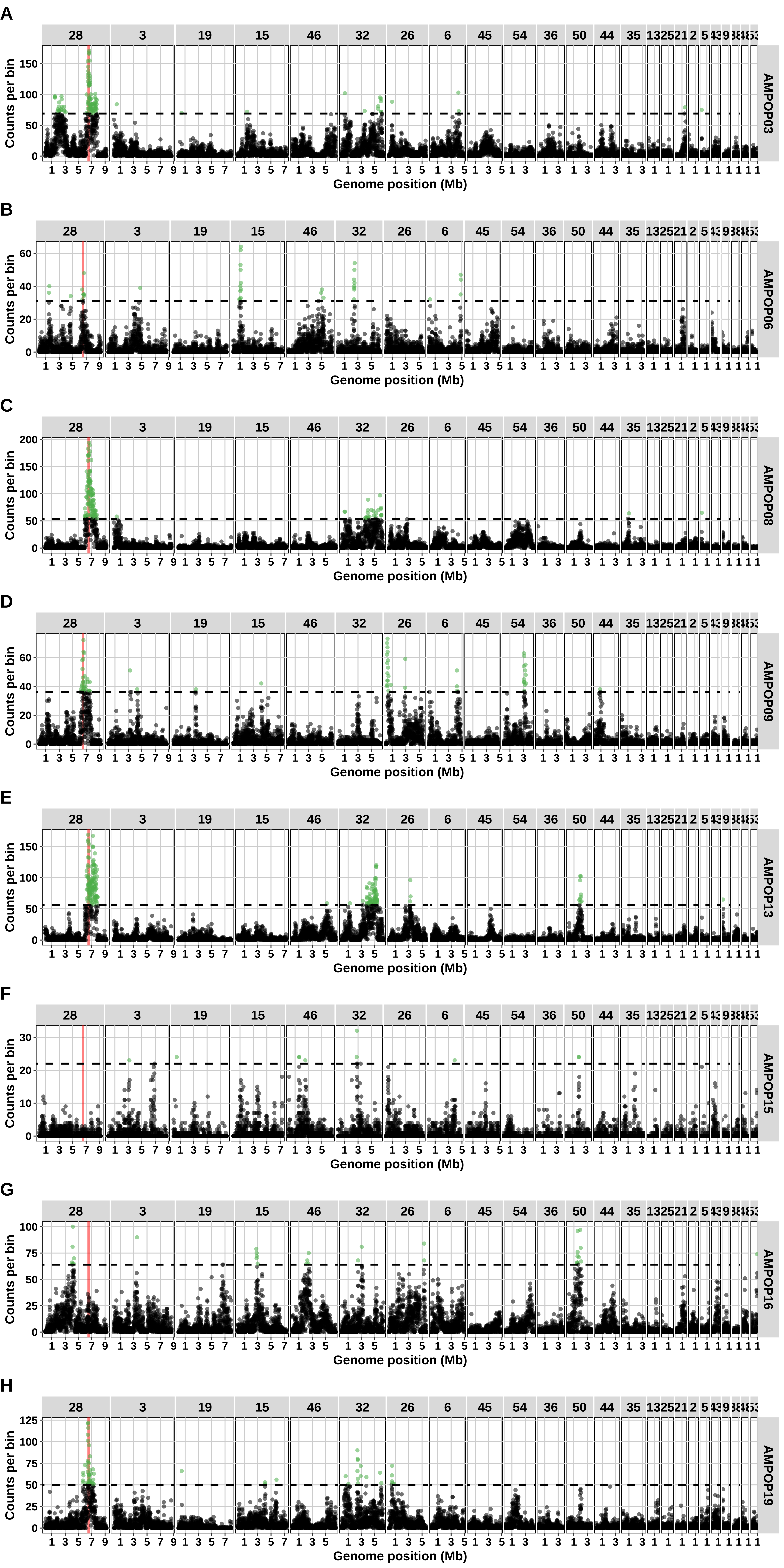

### Supplementary figure 3

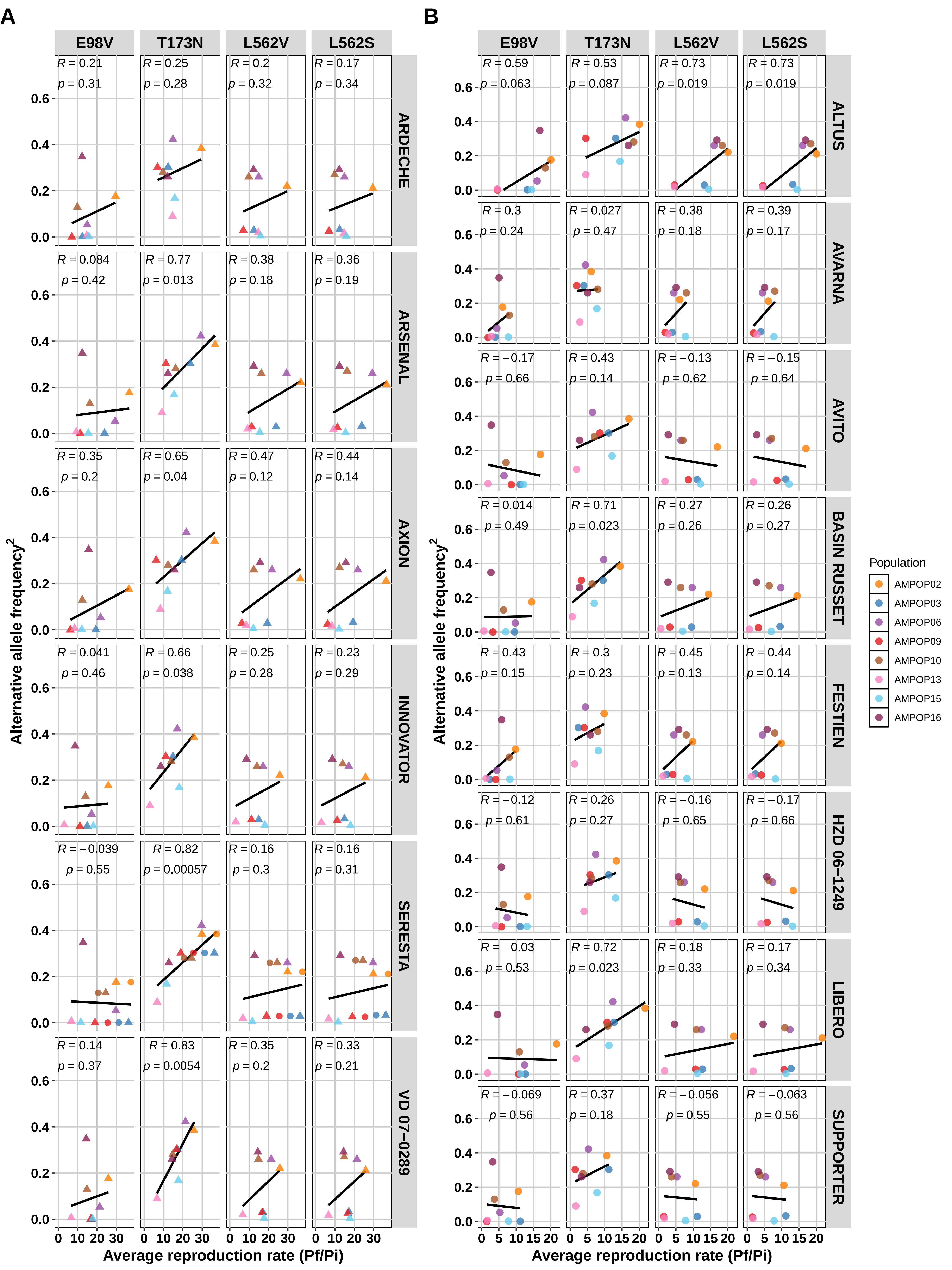

### Supplementary figure 4

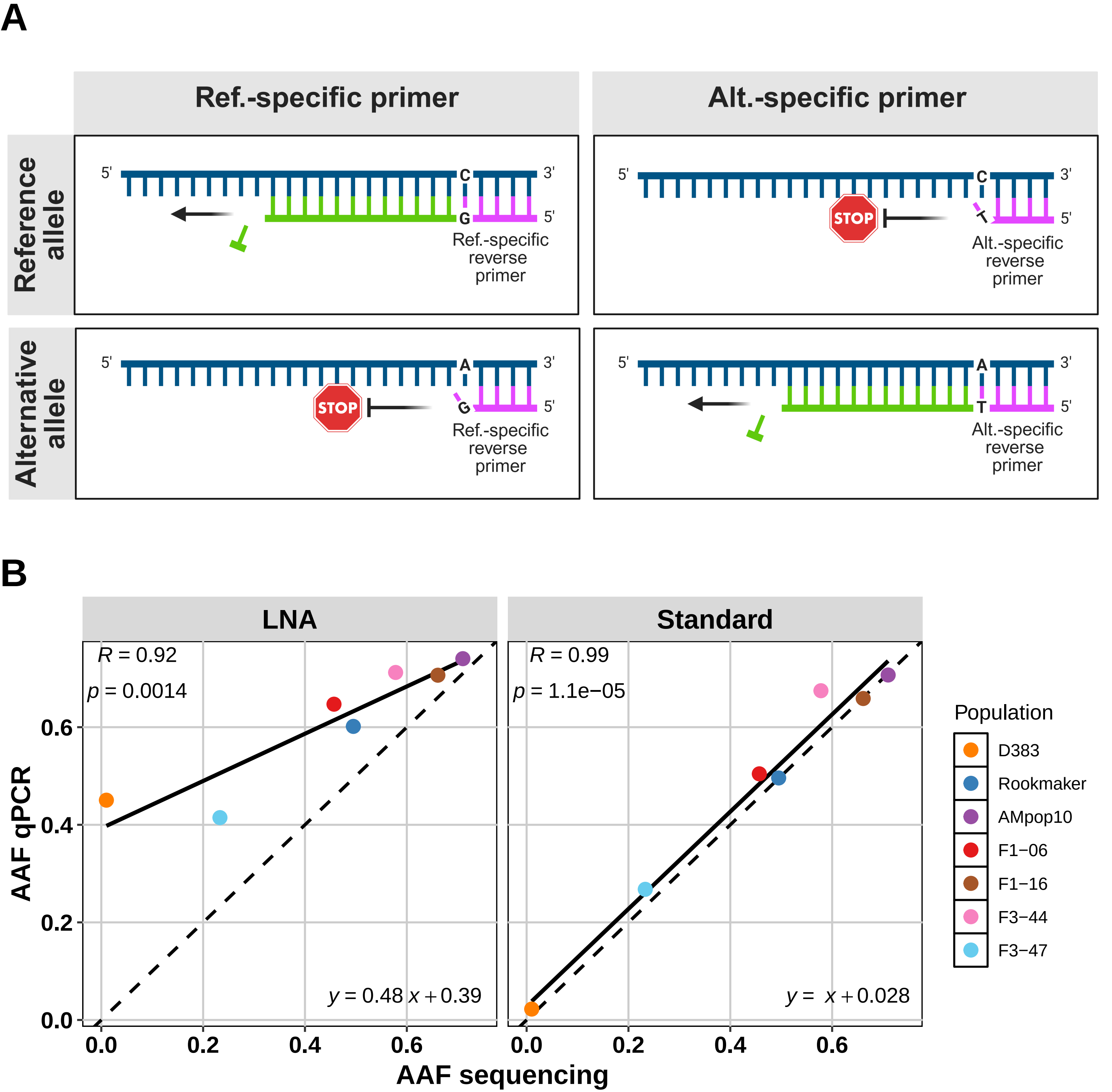

### Supplementary figure 5

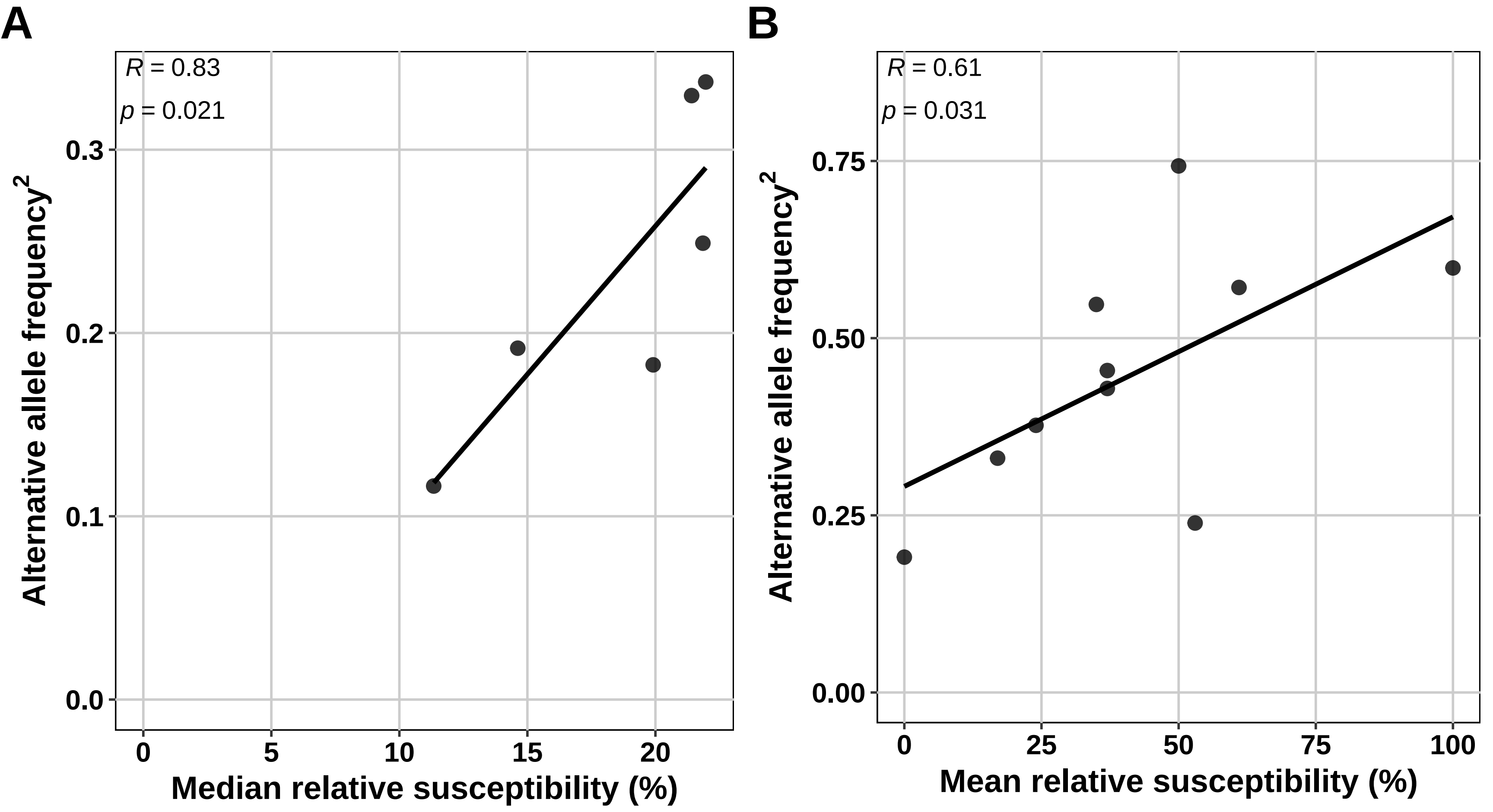

### Supplementary figure 6

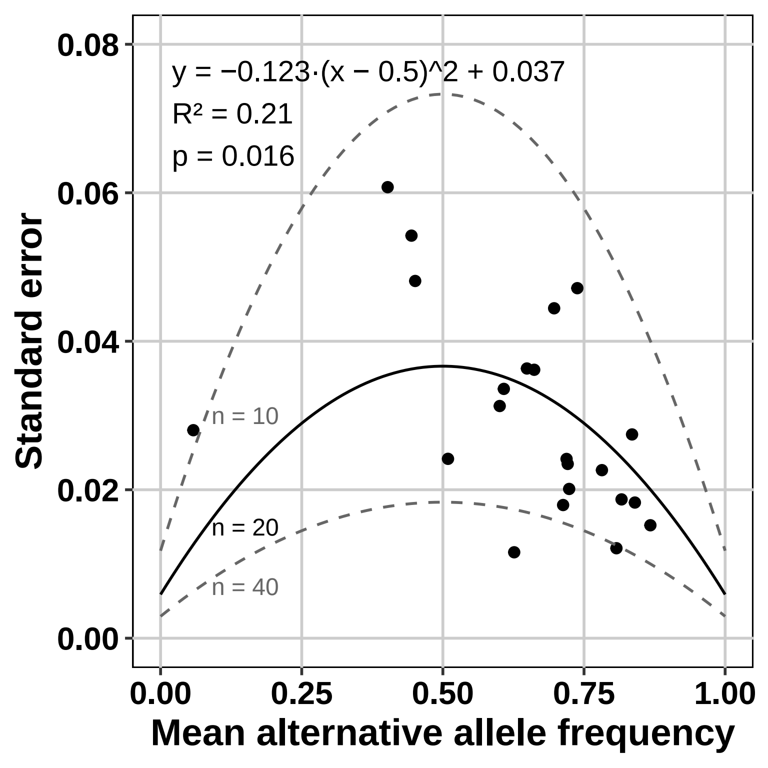
